# Early pathogenesis of spinal and bulbar muscular atrophy uncovered by human iPSC-derived motor neurons highlights pathogenic neuropeptides as therapeutic targets

**DOI:** 10.64898/2025.12.10.692869

**Authors:** Kazunari Onodera, Yuichi Riku, Daisuke Shimojo, Akinobu Ota, Muhammad Irfanur Rashid, Rina Okada, Shinichi Yamaguchi, Shinichiro Yamada, Yoshitaka Hosokawa, Mari Yoshida, Yasushi Iwasaki, Manabu Doyu, Gen Sobue, Masahisa Katsuno, Hideyuki Okano, Yohei Okada

## Abstract

Spinal and bulbar muscular atrophy (SBMA) is a neuromuscular disorder caused by the expansion of the polyglutamine tract in the androgen receptor (AR). Motor neurons (MNs) derived from patient-specific induced pluripotent stem cells (iPSCs) robustl y recapitulated early SBMA phenotypes driven by endogenous mutant AR in the absence of testosterone (dihydrotestosterone) and detectable mutant AR aggregation. Notably, endoplasmic reticulum stress markedly exacerbated SBMA pathology. Cross-species integrative analyses of patient-derived neurons and spinal cords of transgenic mouse models revealed high expression of multiple disease-associated neuropeptides, including urotensin II (UTS2), in patient spinal MNs that was correlated with disease onset and progression in iPSC-derived MNs. Downstream signaling analyses of these neuropeptides revealed convergent molecular pathways whose pharmacological inhibition rescued cellular phenotypes. Together, these results establish a human disease model harboring endogenous mutant AR that closely reproduces early SBMA pathology and provides molecular leads for elucidating disease mechanisms and biomarkers and developing therapeutic targets.

## Introduction

Spinal and bulbar muscular atrophy (SBMA) is an adult-onset, slowly progressive neuromuscular disorder caused by the pathogenic expansion of CAG repeats in the androgen receptor (*AR*) gene. SBMA is characterized by muscle atrophy, fasciculations, bulbar palsy, and facial and proximal muscle weakness^1–4^. Currently, there is no curative treatment for SBMA. To date, the mechanisms underlying SBMA neurodegeneration have been investigated primarily in mouse models, in which mutant AR translocates to the nucleus in a ligand (testosterone)-dependent manner and forms aggregates. In this process, mutant AR exerts cytotoxic effects that eventually lead to neuronal degeneration. Selective neuronal vulnerability has been attributed to transcriptional dysregulation caused by mutant AR aggregation and disruption of proteolytic pathways^4,5^. However, pronounced interspecies differences between human patients and murine models have been reported, leaving the precise disease mechanisms largely unresolved. For example, human patients have been reported to exhibit a pathogenic threshold of 38 or more CAG repeats, whereas mouse models generally require expansions of approximately 100 repeats^6,7^. Human patients exhibit loss of spinal MNs, whereas mouse models exhibit neuronal atrophy without apparent neuronal cell death^2,7^. Nuclear AR aggregates are rarely detected in human skeletal muscle but are prominent in mouse models, where they are accompanied by severe muscle atrophy and rapidly progressive symptoms^7,8^. Moreover, overexpression of wild-type AR in the skeletal muscle of wild-type mice is sufficient to induce a SBMA-like phenotype^9^. Therapeutic responses also diverge. In murine models, castration results in complete phenotype rescue, and treatment with the luteinizing hormone-releasing hormone (LHRH) agonist leuprorelin greatly improves disease manifestations. In human patients, long-term leuprorelin administration reduces the incidence of pneumonia and mortality and has shown beneficial effects, particularly at the early stages of the disease^10^. However, leuprorelin does not completely prevent muscle weakness and is associated with adverse effects such as infertility and skeletal muscle catabolism^7,11,12^. These discrepancies between humans and mice highlight the need for human-derived models expressing endogenous mutant AR that can accurately recapitulate disease phenotypes and therapeutic responses, providing a robust platform for mechanistic pathogenesis studies. Defining early-stage pathology in such models could facilitate the development of biomarkers for early diagnosis and therapeutic strategies for early intervention, ultimately supporting presymptomatic diagnosis and disease prevention.

Here, we established an early-stage disease model using MNs differentiated from SBMA patient-derived induced pluripotent stem cells (iPSCs). This platform provides patient-specific neurons expressing endogenous mutant AR. Although mutant AR aggregates were not detected in SBMA MNs, these cells replicated the molecular alterations and neurodegenerative phenotypes. Transcriptomic analyses revealed that neuropeptides, notably *UTS2*, are associated with early SBMA pathogenesis and have emerged as promising therapeutic targets.

## Results

### SBMA iPSC-derived MNs differentiate and mature in a manner comparable to that of controls

SBMA patient-derived iPSCs (SBMA2E16, 3E10, and 4E5) and control iPSCs (EKN3, TIGE9, and YFE16) were induced to differentiate into MNs using an established protocol^13,14^ (Fig. 1a). MNs were subjected to adherent culture for up to 4 weeks in the presence or absence of 10 nM dihydrotestosterone (DHT), after which their differentiation efficiency was evaluated. After 1 week, immunocytochemical analysis of MN markers revealed approximately 26–43% HB9^+^ or ISL-1^+^ cells, with SBMA iPSC-derived MNs (SBMA MNs) indicating a slight increase in the proportion of HB9^+^ cells compared with control iPSC-derived MNs (control MNs) (Fig. 1b–d and Supplementary Fig. 1a). Quantitative reverse transcriptase polymerase chain reaction (qRT–PCR) revealed no significant differences in *HB9* or *ISL-1* mRNA expression between control and SBMA MNs, except for an increase in *ISL-1* mRNA expression in SBMA MNs in the presence of DHT (Fig. 1e, f). These results indicate that the efficiency of SBMA iPSC differentiation into MNs was not lower than that of control cells.

**Fig. 1.**
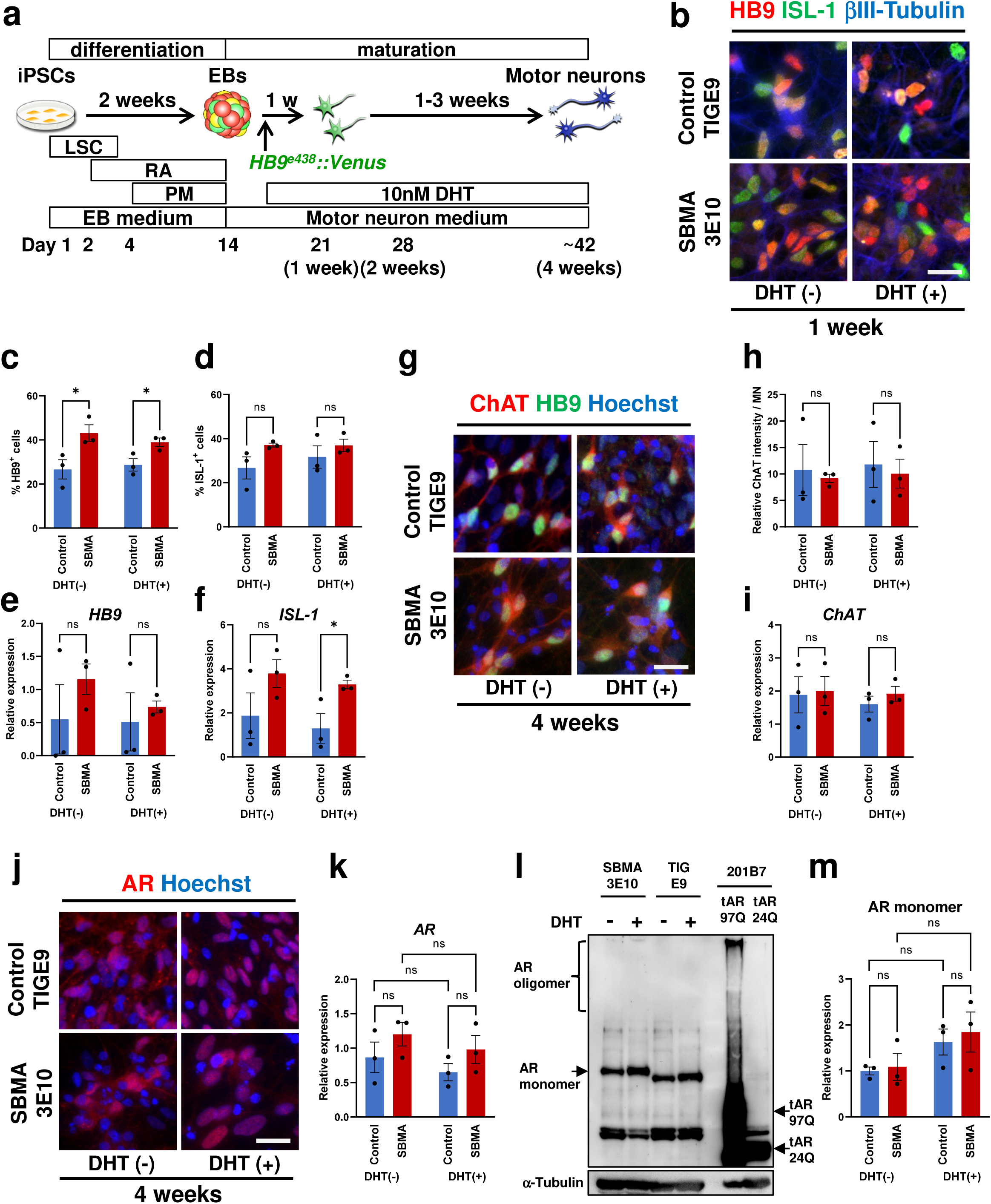
Differentiation and maturation of control and SBMA iPSCs into MNs. **a,** Schematic representation of the differentiation of iPSCs into MNs. LSC, LDN- 193189, SB4315342, and CHIR99021; RA, retinoic acid; PM, purmorphamine. Control clones: EKN3, TIGE9, and YFE16; SBMA clones: SBMA2E16, 3E10, and 4E5. **b,** Immunocytochemical analysis of HB9, ISL-1, and βIII-Tubulin after 1 week of monolayer differentiation with or without 10 nM DHT. Scale bar, 25 μm. **c, d,** Quantification of the HB9^+^ (c) and ISL-1^+^ (d) cells (n = 3). The proportions of HB9^+^ and ISL-1^+^ cells in the SBMA group were no lower than those in the control group. **e, f** Quantitative RT–PCR analysis of *HB9* (**e**) and *ISL-1* (**f**) expression after 1 week of monolayer culture (n = 3). **g,** Immunocytochemical analysis of ChAT expression after 4 weeks of monolayer differentiation with or without 10 nM DHT. Scale bar, 25 μm. **h,** Quantification of ChAT^+^ MN intensity (n = 3). No significant difference in signal intensity was detected between the control and SBMA groups. **i,** Quantitative RT–PCR analysis of *ChAT* (**h**) expression after 4 weeks of monolayer culture (n = 3). **j,** Immunocytochemical analysis of AR expression after 4 weeks of monolayer differentiation. Scale bar, 25 μm. DHT-dependent nuclear translocation of AR was observed, but nuclear AR aggregation was not detected. **k,** Quantitative RT–PCR analysis of AR expression after 4 weeks of monolayer differentiation (n =3). No significant difference was observed between the control and SBMA MNs. **l,** Western blot analysis of AR protein expression after 4 weeks of monolayer differentiation. No AR oligomers were observed in SBMA MNs treated with 10 nM DHT. As a positive control, aggregated oligomeric AR was observed in 201B7 MNs expressing truncated AR-97Q (tAR-97Q), whereas no AR aggregates were detected in 201B7 MNs expressing tAR24Q. See also Supplementary Fig. 1e. **m,** Quantification of the AR monomer expression in (**l**). Treatment with DHT increased the level of AR monomers. See also Supplementary Fig. 1e. Data are presented as the mean ± SEM; *, *p* < 0.05; **, *p* < 0.01; Student’s *t* test.

To assess MN maturation, choline acetyltransferase (ChAT) expression was assessed by immunocytochemistry, Western blotting, and qRT–PCR after 4 weeks of differentiation. No significant differences in ChAT expression were detected between control and SBMA MNs in any of these analyses, suggesting that SBMA MNs did not exhibit impaired maturation (Fig. 1g–i and Supplementary Fig. 1c, d).

### SBMA-iPSC-derived MNs recapitulate early disease pathology prior to mutant AR aggregation

AR dynamics were examined in MNs after 4 weeks of differentiation and immunostaining revealed no obvious differences between control and SBMA MNs. In both types of MNs, DHT treatment induced the nuclear translocation of AR and mutant AR (Fig. 1j and Supplementary Fig. 1b). qRT–PCR analysis revealed no significant changes in *AR* mRNA expression upon DHT treatment (Fig. 1k), whereas Western blotting revealed a trend toward a DHT-dependent increase in AR monomer levels (Fig. 1l, m and Supplementary Fig. 1e). Furthermore, mutant AR aggregates were not observed in SBMA MNs by either immunocytochemistry or Western blotting, even after prolonged exposure time (Fig. 1j, l and Supplementary Fig. 1e). These findings indicate that SBMA MNs exhibited no overt abnormalities, including mutant AR aggregates, during the 4-week differentiation period.

To assess neurite integrity, MNs were transduced with *HB9^e438^::Venus* lentivirus, a MN-specific reporter, by which Venus fluorescent protein, a variant of yellow fluorescent protein (YFP)^15^, was expressed under the control of the HB9^e438^ enhancer and β-globin minimal promoter on day 2 of differentiation, which coincided with the initiation of DHT treatment^13^. Neurite length in the presence or absence of DHT was then monitored for 8 weeks. Progressive neurite degeneration was observed in SBMA MNs beginning at 4 weeks, and by 8 weeks, SBMA MN neurite length was reduced by 34% compared with control MNs (Fig. 2a, b). DHT treatment did not significantly influence neurite outgrowth in SBMA MNs.

**Fig. 2.**
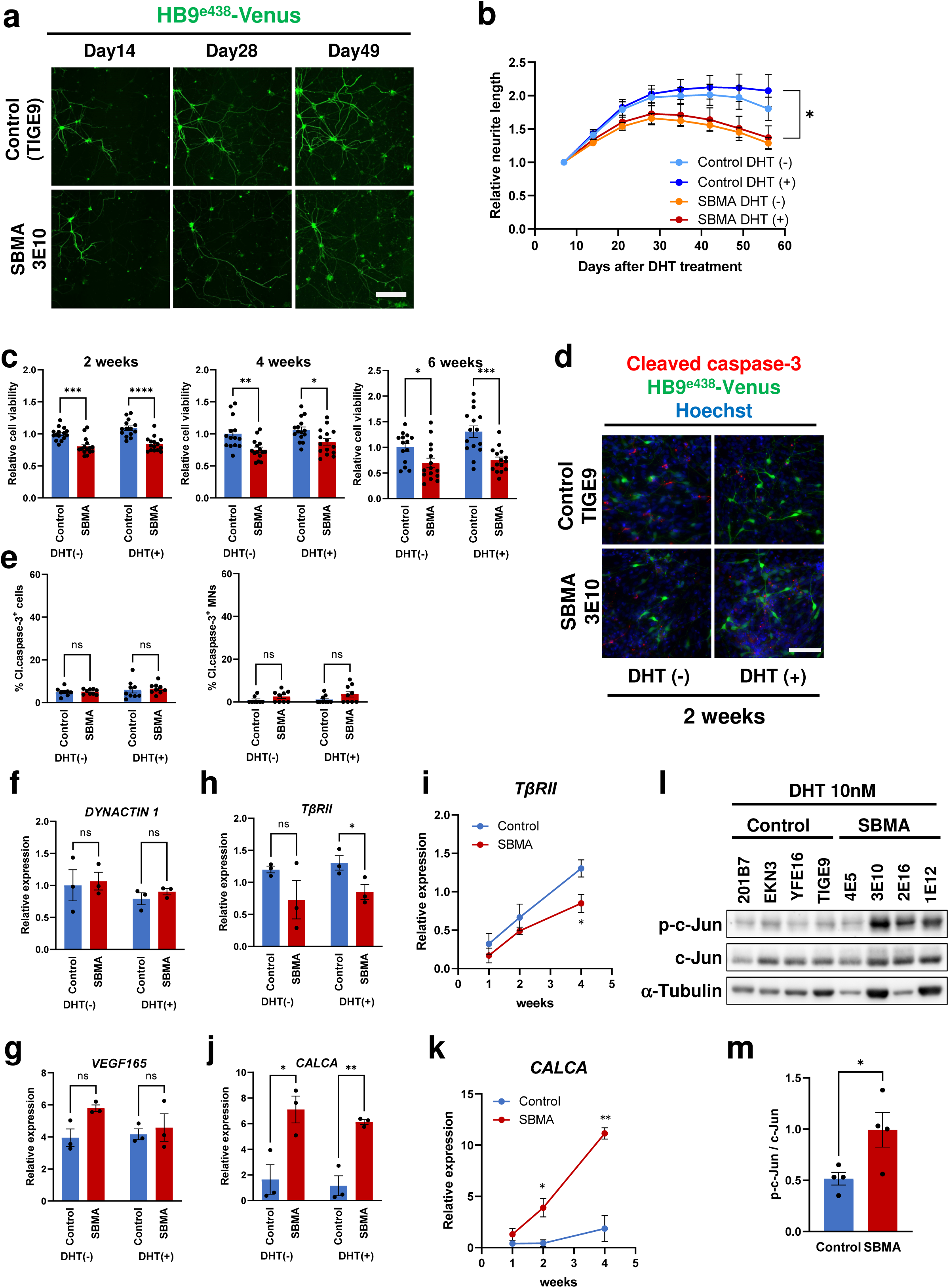
SBMA iPSC-derived MNs recapitulated early SBMA pathology. **a,** Representative live-cell images of neurites in the presence of 10 nM DHT. Scale bar, 100 μm. **b,** Neurite length 8 weeks after DHT addition (n = 9). After 3 weeks, the neurite length of SBMA MNs gradually decreased with or without DHT treatment. **c,** Cell viability was assessed by the MTS assay (n = 3). The cell viability of the SBMA MNs progressively decreased over time. **d,** Immunocytochemical analysis of cleaved caspase-3 expression after 2 weeks of monolayer differentiation. Scale bar, 100 μm. **e,** Quantification of the proportion of cleaved caspase-3^+^ cells among all cells (left) and among all MNs (right) (n = 3). No significant differences were detected between the control and SBMA MNs. **f, g,** Quantitative RT–PCR analysis of *DYNACTIN1* (**f**) and *VEGF165* (**g**) expression after 4 weeks of monolayer differentiation (n = 3). No significant differences were detected between the control and SBMA MNs. **h, i,** Quantitative RT–PCR analysis of *TβRII* expression after 4 weeks of monolayer differentiation (**h**) and time course analysis (**i**). *TβRII* expression was significantly lower in SBMA MNs than in control MNs at 4 weeks. **j, k,** Quantitative RT–PCR analysis of *CALCA* expression after 4 weeks of monolayer differentiation (**j**) and time course analysis (**k**). *CALCA* expression was significantly higher in SBMA MNs than in control MNs after two and 4 weeks of monolayer differentiation. **l, m,** Western blot analysis of c-Jun expression after 1 week of monolayer differentiation. Phospho-c-Jun (p-c-Jun) expression was significantly higher in SBMA MNs (SBMA1E12, 2E16, 3E10, and 4E5) than in control MNs (201B7, EKN3, TIGE9, and YFE16). Quantification of p-c-Jun expression is shown in (**m**) (n = 4). The data are presented as the mean ± SEM; *, *p* < 0.05; **, *p* < 0.01; Student’s *t* test; one-way ANOVA; or two-way repeated-measures ANOVA followed by the Tukey–Kramer multiple comparisons test.

Cell viability was evaluated via the use of the MTS assay, which reflects mitochondrial dehydrogenase activity. In SBMA MNs, MTS activity was significantly reduced after 2 weeks both in the presence and absence of DHT and progressively decreased, reaching approximately 43% (with DHT) and 38% (without DHT) that of control MNs by 6 weeks. However, no clear dependency on DHT was observed (Fig. 2c). Despite the reduced MTS activity, lactate dehydrogenase (LDH) leakage did not increase in SBMA MNs compared with that in controls, and the proportion of cleaved caspase-3^+^ MNs labeled by HB9-Venus fluorescence was not significantly increased according to the immunocytochemistry results (Fig. 2d, e and Supplementary Fig. 2a, b). These data suggest that SBMA MNs exhibit mitochondrial dysfunction without progression to overt neuronal death^16,17^.

To investigate molecular changes in SBMA MNs, we performed qRT–PCR analysis for genes previously implicated in SBMA. No significant differences were detected in the expression of *DYNACTIN1*, which regulates retrograde axonal transport, or *VEGF165*, which supports MN survival, although both were reported to be reduced in the context of SBMA (Fig. 2f, g). In contrast, the level of *transforming growth factor-*β *receptor type II* (*T*β*RII*), which mediates neuroprotective TGFβ signaling, began to decrease at 2 weeks and was significantly reduced by 4 weeks in SBMA MNs compared with that in controls, which is consistent with the findings of previous reports (0.65 ± 0.09-fold that of the control, n = 3, *p* < 0.05) (Fig. 2h, i)^5^. Notably, the expression of *Calcitonin- related polypeptide alpha* (*CALCA*), encoding calcitonin gene-related peptide 1 (CGRP-1), was significantly upregulated in SBMA MNs at 2 weeks and further increased to 5.31 ± 0.17-fold that of the control by 4 weeks (n = 3, *p* < 0.01) (Fig. 2j, k). Given that CGRP-1 has been reported to induce neuronal toxicity via c-Jun phosphorylation within the context of SBMA, we examined c-Jun phosphorylation after 1 week of DHT treatment^18^. Western blotting revealed approximately 1.9-fold higher levels of phospho-c-Jun in SBMA MNs than in controls (n = 4, *p* < 0.05), which is consistent with previous findings, but clear dependency on DHT was not observed (Fig. 2l, m).

Overall, despite the absence of apparent AR aggregation or neuronal death, SBMA MNs exhibited intrinsic transcriptional dysregulation, mitochondrial dysfunction, and progressive neurodegenerative features such as decreased neurite integrity, thereby faithfully recapitulating the early-stage pathology that precedes mutant AR aggregation.

### ER stress exacerbates SBMA disease progression

We next investigated how cellular stress, which has been implicated in aging, contributes to disease progression in the context of SBMA^19^. Control and SBMA MNs subjected to 4 weeks of differentiation were exposed to various stress inducers targeting autophagy, the proteasome, endoplasmic reticulum (ER) stress, oxidative stress, and other pathways for 72 hours. Polyglutamine (polyQ) signals were then examined by Western blotting using 1C2, an antibody that specifically recognizes expanded polyQ tracts. Among these treatments, tunicamycin (TM) induced more than a 10-fold increase in the polyQ signal in SBMA MNs (Supplementary Fig. 3a, b). Because TM activates ER stress by blocking N-linked glycosylation of nascent proteins and because ER stress is associated with increased levels of mutant AR protein^20^, which exacerbates SBMA pathology, we subsequently focused on ER stress.

To assess the basal ER stress status, we measured the expression levels of genes associated with the unfolded protein response (UPR) in control and SBMA MNs cultured for 4 weeks without ER stress inducers by qRT–PCR. No significant differences in the expression of *binding immunoglobulin protein* (*BIP*), *C/EBP homologous protein* (*CHOP*), *spliced X-box binding protein 1* (*sXBP1*), *activating transcription factor 4* (*ATF4*), *HMG-CoA reductase degradation 1* (*HRD1*), or *ER degradation-enhancing alpha-mannosidase-like protein* (*EDEM*) were detected between the two groups (Supplementary Fig. 3c). In contrast, Western blotting at 4 weeks revealed altered expression of ER stress-related proteins in SBMA MNs: the expression of protein disulfide isomerase (PDI) was increased (2.98 ± 0.12-fold), whereas that of eukaryotic translation initiation factor 2 alpha (eIF2α) and ATF6 decreased (0.80 ± 0.04-and 0.49 ± 0.10-fold, respectively) (Fig. 3a, b). Moreover, exposure of SBMA MNs to thapsigargin (TG; 200 nM), a selective ER Ca^2+^-ATPase inhibitor, for 4–48 hours significantly upregulated the expression of *CHOP*, *sXBP1*, and *ATF4* (Fig. 3d). These changes are consistent with the increased vulnerability to ER stress associated with the expression of mutant AR.

**Fig. 3.**
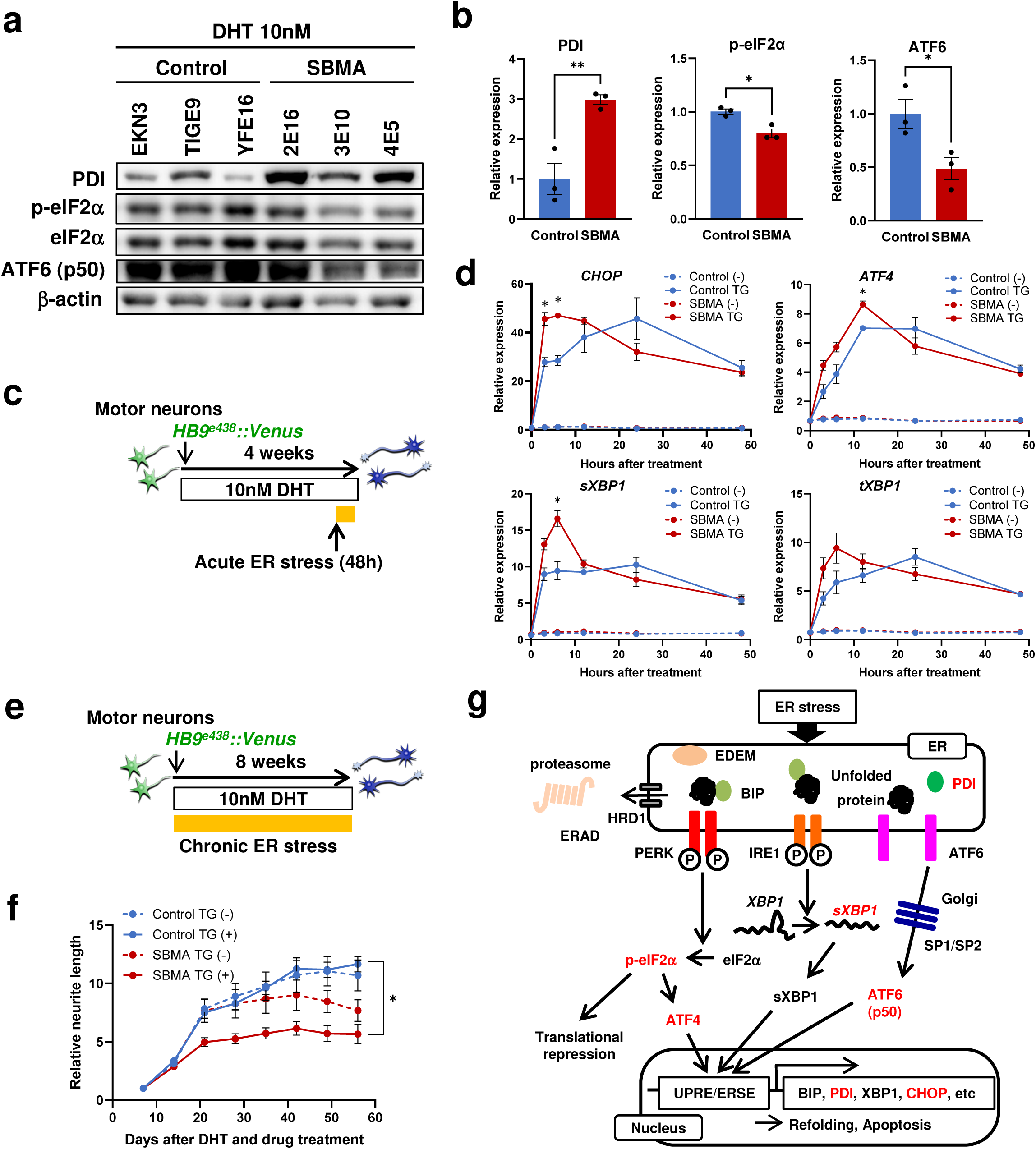
SBMA MNs are vulnerable to ER stress. **a,** Western blot analysis of the expression of ER stress-related proteins. **b,** Quantification of the expression levels of ER stress-related proteins shown in (**a**) (n = 3). **c,** Schematic representation of the analysis of acute ER stress after 4 weeks of differentiation. **d,** Time course qRT–PCR analysis of genes related to the unfolded protein response (n = 3) (**d**). *CHOP*, *ATF4*, and *sXBP1* expression was significantly upregulated in SBMA MNs within 6 hours of treatment with 200 nM thapsigargin (TG). **e,** Schematic representation of the analysis of chronic ER stress during 8 weeks of monolayer differentiation. **f,** Relative neurite length of control and SBMA MNs in the presence of 1 nM TG for 8 weeks (n = 4). A significant decrease in neurite length was observed in SBMA MNs after 3 weeks of treatment with TG. Control iPSC clones: EKN3, TIGE22, YFE16, and YFE19; SBMA iPSC clones: SBMA1E12, 2E16, 3E10, and 4E5. **g,** Schematic representation of ER stress signaling pathways. Dysregulation of the PERK and IRE1 branches of the UPR was observed in SBMA MNs. Data are presented as the mean ± SEM; *, *p* < 0.05; **, *p* < 0.01; Student’s *t* test; one-way ANOVA; or two-way repeated-measures ANOVA followed by the Tukey–Kramer multiple comparisons test.

To confirm the long-term effects of ER stress on SBMA MN pathology, control and SBMA MNs were maintained for up to 8 weeks in the presence or absence of a low concentration of TG (1 nM) to avoid the acute cytotoxicity induced at high concentrations (Fig. 3e). After 3 weeks of TG exposure, the SBMA MN neurite length was reduced to 0.65 ± 0.08-fold that of untreated MNs, whereas control MN neurite length was unaffected (Fig. 3f). These results indicate that chronic ER stress accelerates neurite degeneration and exacerbates SBMA-associated phenotypes.

Finally, we analyzed which branches of the UPR mediate disease progression in SBMA MNs. The UPR comprises three major signaling branches activated by the accumulation of unfolded or misfolded proteins in the ER: the inositol-requiring enzyme 1 (IRE1) branch, the ATF6 branch, and the protein kinase R-like ER kinase (PERK) branch. Our findings indicate that both the IRE1 branch, as evidenced by increased PDI and sXBP1 expression, and the PERK branch, as evidenced by elevated CHOP and ATF4 expression, were activated (Fig. 3g). Taken together, these findings suggest that SBMA MNs are particularly vulnerable to ER stress through the concurrent activation of the IRE1 and PERK branches of the UPR.

### *CALCA*, *UCN, UTSII*, and *VIP* are upregulated in SBMA MNs

To identify the molecular mechanisms underlying SBMA pathogenesis, we performed transcriptome analyses of iPSC-derived MNs comprising two pairs of control (TIGE9) and SBMA (3E10) MNs cultured for 4 weeks. This analysis revealed 2,394 upregulated genes and 1,562 downregulated genes whose expression changed more than 1.5-fold (Supplementary Tables 2 and 3). Via the use of a previously reported dataset of spinal cord samples from mutant AR transgenic mice (AR-24Q and AR-97Q mice) at preclinical, early, and advanced disease stages, we also identified differentially expressed genes (DEGs) whose expression was significantly altered between AR-24Q and AR-97Q mice at any stage^18^. This yielded 316 DEGs in the preclinical stage, 138 at the early stage, and 164 in the advanced stage (Supplementary Table 1).

We subsequently identified the genes that overlapped between the mouse spinal cord and human iPSC-MN datasets, resulting in the identification of 46 upregulated and 30 downregulated genes (Supplementary Fig. 4a and Supplementary Table 4). Notably, approximately 78% of these genes corresponded to the preclinical and early stages, supporting that SBMA MNs recapitulate early pathological changes (Fig. 4a).

**Fig. 4.**
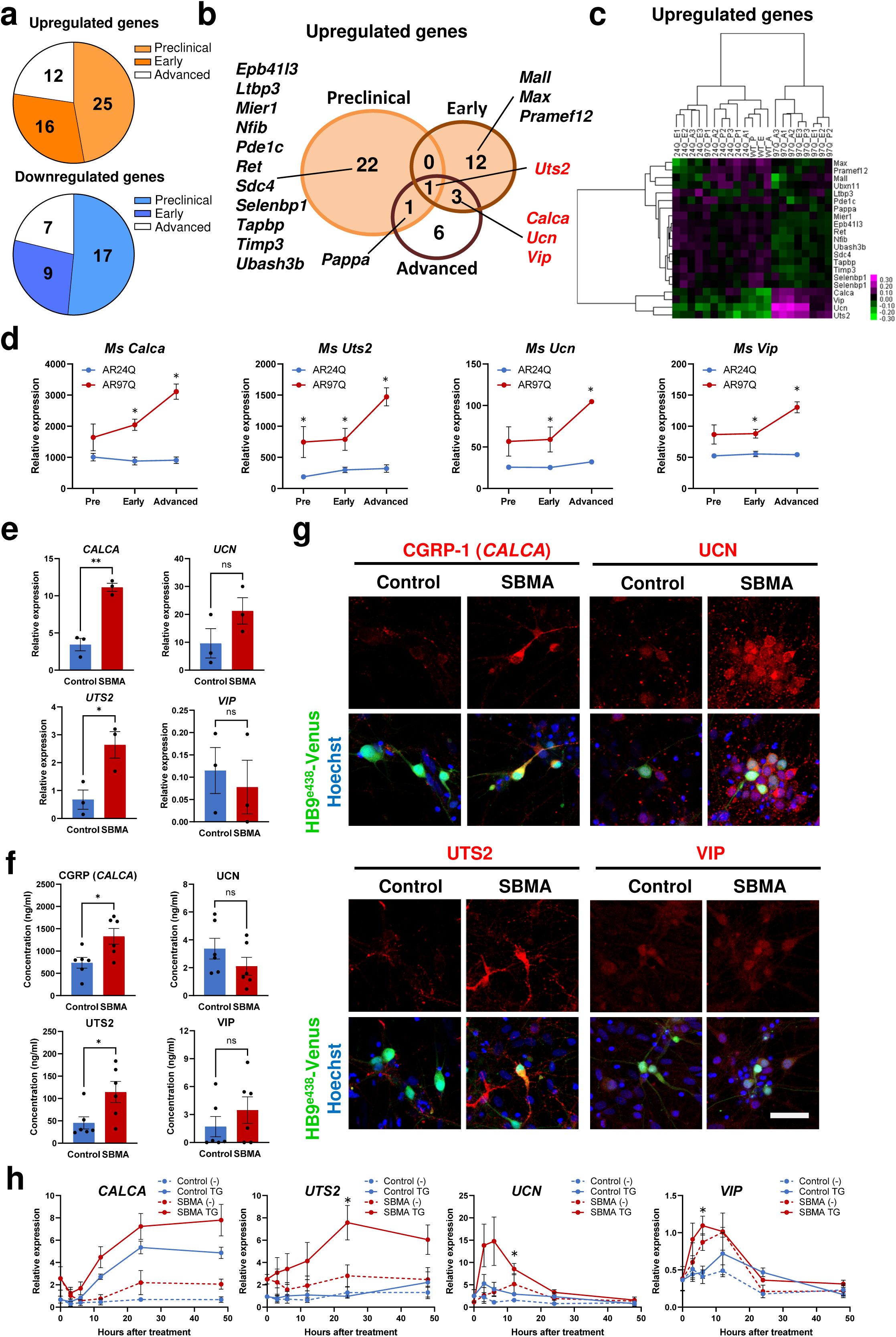
*CALCA*, *UCN*, *UTS2*, and *VIP* are upregulated in SBMA MNs. **a,** Number of genes overlapping between the human and mouse microarray analyses. Among a total of 86 genes, 78% of commonly altered genes were detected at the preclinical and early stages. SBMA MNs closely replicated early pathology. **b,** Venn diagram showing genes commonly upregulated in human SBMA patients and model mice at the preclinical, early, and advanced stages. **c,** Hierarchical clustering analysis of upregulated genes. The rows in the heatmap represent selected genes, and the columns represent the mouse groups. P, preclinical; E, early; A, advanced; WT, wild type; 24Q, AR-24Q; and 97Q, AR-97Q. **d,** Time course of *Calca*, *Ucn*, *Uts2*, and *Vip* expression in mice. The expression of all four genes gradually increased over time (n = 3). **e,** Quantitative RT–PCR analysis of *CALCA, UCN, UTS2* and *VIP* expression in control and SBMA iPSC-derived MNs (SBMA MNs) after 4 weeks of monolayer differentiation (n = 3). The expression of *CALCA* and *UTS2* was significantly greater in SBMA MNs than in control MNs. **f,** ELISA of culture supernatants after 4 weeks of MN culture (n = 3). CGRP and UTS2 protein levels were increased in SBMA MNs. **g,** Immunocytochemical analysis of CGRP-1, UCN, UTS2 and VIP expression in control and SBMA MNs after 4 weeks of differentiation. Scale bar, 50 μm. All four proteins were detected. **h,** Time course analysis of *CALCA, UCN, UTS2* and *VIP* expression under ER stress by qRT–PCR (n = 3). After 4 weeks of monolayer differentiation, the MNs were treated with 200 nM TG for 48 hours. TG treatment increased the expression of all four genes in SBMA MNs but not in control MNs. Data are presented as the mean ± SEM; *, *p* < 0.05; **, *p* < 0.01; Student’s *t* test; one-way ANOVA; or two-way repeated-measures ANOVA followed by the Tukey–Kramer multiple comparisons test. In (d), a moderated *t* test with a Westfall–Young permutation multiple comparison test was used.

After removing probe-level duplicates, we further identified 19 upregulated and 14 downregulated genes that were particularly prominent in the preclinical and early stages among the genes that exhibited stage-dependent dynamic changes in expression during disease progression (Fig. 4b, c and Supplementary Fig. 4b, c). Hierarchical clustering revealed 4 upregulated genes as early disease-related markers, namely, *CALCA*, *urocortin* (*UCN*), *urotensin II* (*UTS2*), and *vasoactive intestinal peptide* (*VIP*), each of which encode a neuropeptide (Fig. 4c). These 4 genes were also upregulated in AR-97Q SBMA mice compared with in AR-24Q mice from the preclinical stage, and their expression further increased with disease progression (Fig. 4d).

To validate these early disease-related markers, we examined their expression in iPSC-derived MNs. qRT–PCR demonstrated that the mRNA levels of *CALCA* (3.24 ± 0.16-fold) and *UTS2* (3.90 ± 0.70-fold) were significantly greater in SBMA MNs than in control MNs at 4 weeks, whereas the expression of *UCN* and *VIP* did not significantly differ between the groups (n = 3, *p* < 0.05) (Fig. 4e). At the protein level, neuropeptide secretion into the culture supernatants measured by enzyme-linked immunosorbent assay (ELISA) was consistent with the transcript data: CGRP (*CALCA*) and UTS2 levels were increased in SBMA MNs, whereas UCN and VIP levels remained unchanged (Fig. 4f). Immunostaining confirmed the robust expression of CGRP, UCN, and UTS2, while VIP expression was low in both control and SBMA MNs (Fig. 4g).

Given the increased vulnerability of SBMA MNs to ER stress, we also examined how ER stress affects the expression dynamics of these early disease-related markers. *CALCA* and *UTS2* expression progressively increased over 24 hours, whereas *UCN* and *VIP* expression transiently increased at 6–12 hours before returning to baseline levels (Fig. 4h). These results demonstrate that ER stress, which promotes disease progression, increases the expression of early pathology-associated markers, further supporting the involvement of these markers at the early stages of SBMA pathology.

### The expression of the identified neuropeptides is confirmed in postmortem spinal cord tissues from SBMA patients

To validate the *in vivo* expression of the four neuropeptides (CGRP, UTS2, UCN, and VIP), we examined their expression in postmortem spinal cord samples from 3 SBMA patients and 2 male control subjects without neurological disorders using immunohistochemistry. The demographic characteristics are summarized in Supplementary Table 5, and all the SBMA patients had expanded CAG repeats, whereas all the control subjects had normal CAG repeat lengths. The expression of CGRP, UTS2, and UCN was significantly greater in the spinal cord MNs of SBMA patients than in those of controls, whereas VIP expression was undetectable in both control and patient MNs (Fig. 5 and Supplementary Fig. 5a).

**Fig. 5.**
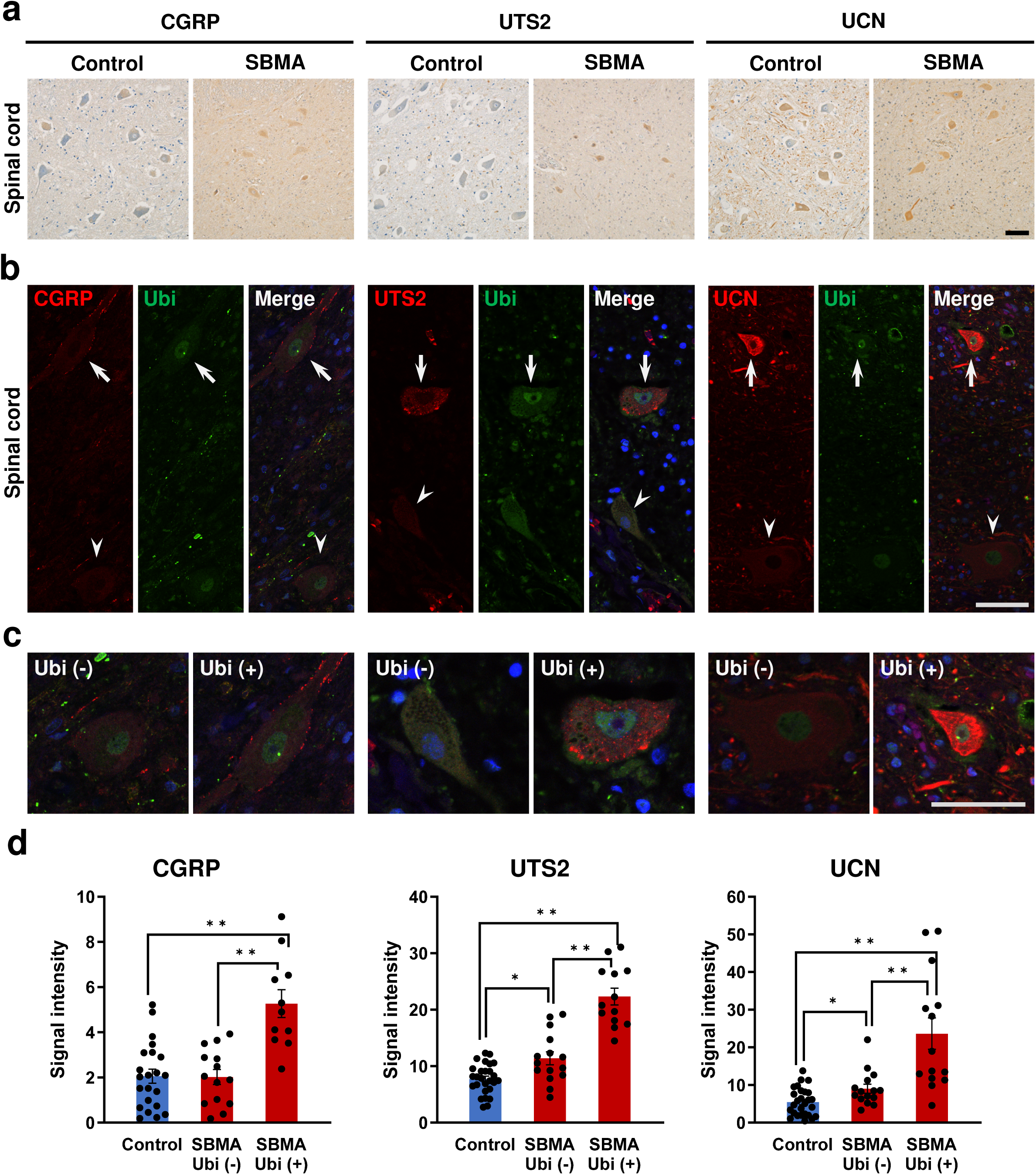
CGRP, UTS2, and UCN are more abundantly expressed in the spinal cords of SBMA patients. **a,** Immunohistochemical analysis of the anterior horn of lumbar spinal cords from SBMA patients and healthy controls. Spinal MNs from SBMA patients showed immunoreactivity for CGRP, UTS2, and UCN. In contrast, control MNs were largely immunonegative for CGRP and UTS2, while a subset exhibited UCN immunoreactivity. Scale bar, 50 μm. **b,** Double immunostaining for ubiquitin (Ubi) and neuropeptides in SBMA spinal cords. Neurons with intranuclear ubiquitin inclusions (NIs, arrows) exhibited higher signal intensities of CGRP, UTS2, and UCN than neurons lacking inclusions did (arrowheads). Scale bar, 20 μm. **c,** Higher magnification images of the neurons indicated by the arrows or arrowheads in (**b**). Scale bar, 20 μm. **d,** Quantification of CGRP, UTS2, and UCN expression in MNs with or without ubiquitin^+^ NIs. MNs with ubiquitin^+^ NIs showed significantly higher neuropeptide signal intensities than NI-negative or control MNs did (Kruskal–Wallis test; *, *p* < 0.05; **, *p* < 0.01).

A subset of SBMA MNs form ubiquitin-positive intranuclear inclusions (NIs), which exhibit granular or diffuse intranuclear immunoreactivity and are thought to express mutant AR^21^. We therefore measured the fluorescence intensities of CGRP, UTS2, and UCN in SBMA spinal cord MNs with and without NIs for comparison with control spinal cord MNs. The fluorescence intensity of each of the three early disease-related neuropeptides was significantly greater in the MNs with NIs than in the MNs without NIs and in the controls (Fig. 5b–d). Together, these findings suggest that the neuropeptides CGRP, UTS2, and UCN may contribute to the pathogenesis of SBMA.

### SBMA pathology is recapitulated by neuropeptide overexpression and rescued by neuropeptide knockdown

To determine whether the four neuropeptides identified in SBMA MNs contribute to disease pathogenesis, control MNs were transduced with lentiviral vectors for the expression of each neuropeptide (UTS2, CGRP-1, UCN, and VIP) along with Venus fluorescent protein under the control of the HB9^e438^ enhancer and β-globin minimal promoter (*HB9^e438^::Gene of Interest (GOI)-IRES-Venus*), and their phenotypes were examined (Fig. 6a and Supplementary Fig. 6a, b). Forced expression of each neuropeptide significantly reduced neurite outgrowth by approximately 20–30%, resulting in neurite lengths comparable to those observed in SBMA MNs (n = 5, *p* < 0.05; Fig. 6b and Supplementary Fig. 6a, b). Because *CGRP-1* and *CALCITONIN* (*CTN*) are splice variants of *CALCA*, we additionally examined *CTN* overexpression, which similarly suppressed neurite outgrowth (n = 5, *p* < 0.05; Fig. 6b).

**Fig. 6.**
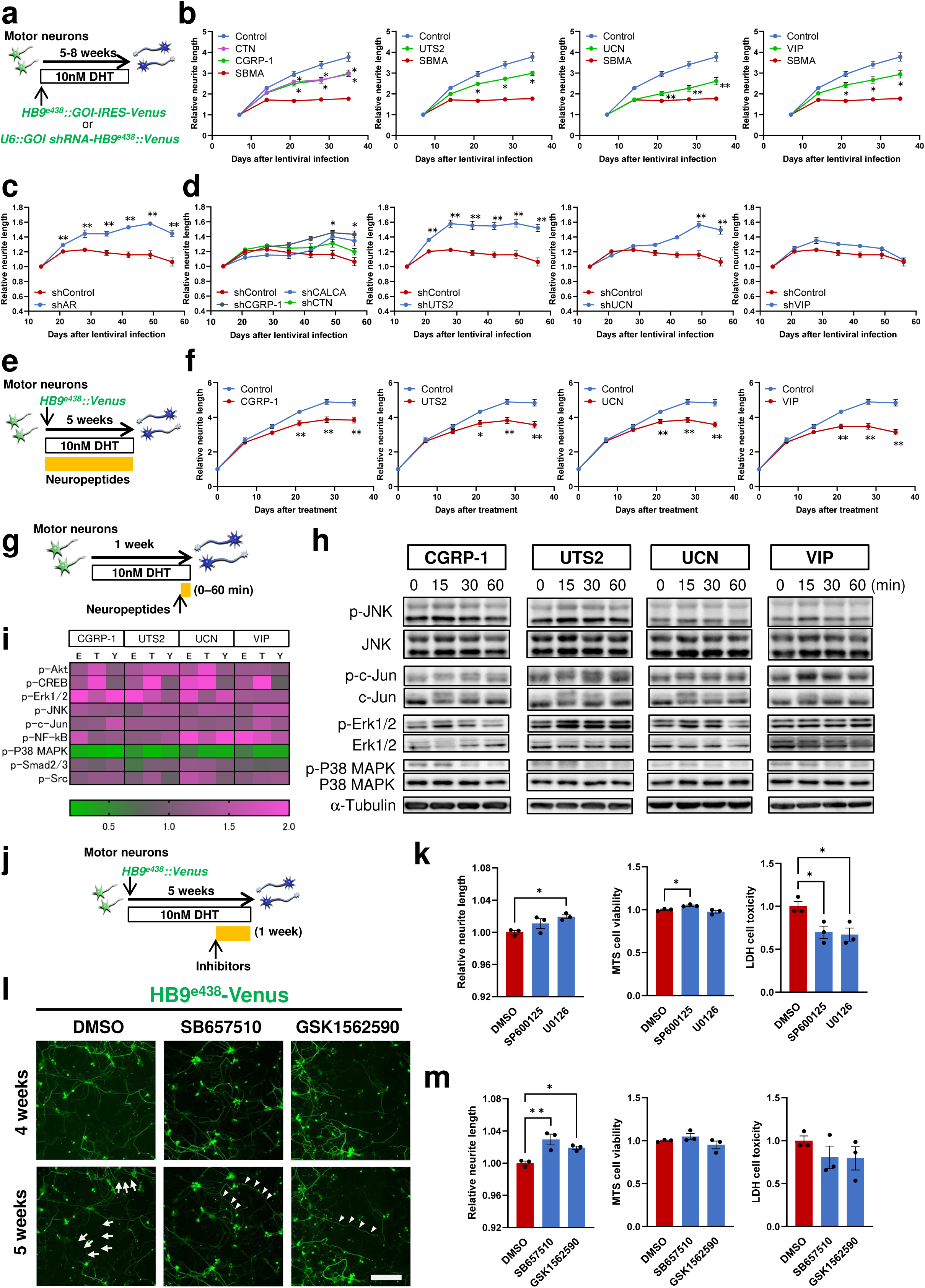
JNK, ERK1/2, and Urotensin II receptor inhibitors alleviate MN degeneration. **a,** Schematic representation of the overexpression of the four neuropeptides in control MNs (YFE16) or their knockdown in SBMA MNs (SBMA3E10). **b,** Neurite lengths of control MNs (YFE16) overexpressing *CGRP-1, CALCITONIN (CTN), UTS2, UCN*, or *VIP*. Overexpression induced significant neurite degeneration after day 11 (n = 6). **c, d,** Neurite lengths of SBMA MNs (SBMA3E10) transduced with lentivirus expressing shRNA targeting AR or neuropeptides or control shRNA. Rescue of neurite outgrowth was observed with shRNAs targeting *AR* (**c**), *CALCA, CGRP-1, CTN, UTS2*, and *UCN* (**d**) (n = 3 for shRNA for neuropeptides; n = 9 for control shRNA). **e,** Schematic representation of the treatment of control MNs (EKN3, TIGE9, and YFE16) with each neuropeptide (CGRP-1, UTS2, UCN, and VIP). **f**, Neurite lengths of control MNs treated with 1 nM of each neuropeptide. Significant neurite degeneration was observed after 3 weeks after treatment with each of the neuropeptides (n = 3). **g,** Schematic representation of the treatment of control MNs with each neuropeptide (CGRP-1, UTS2, UCN, or VIP) after 1 week of monolayer culture for the analysis of downstream signals. **h,** Time-course analysis of the expression of phosphorylated signaling proteins in control MNs (TIGE9) treated with each neuropeptide by Western blotting. See also Supplementary Fig. 6f. **i,** Heatmap of the expression of phosphorylated signaling proteins. E, EKN3; T, TIGE9; and Y, YFE16. **j,** Schematic representation of the treatment of SBMA MNs with signaling inhibitors or UTS2 receptor (UTS2R) inhibitors. **k,** Neurite length (left), cell viability (MTS assay, middle), and cytotoxicity (LDH assay, right) of SBMA MNs treated for 1 week with 0.0001% DMSO or signaling inhibitors (100 nM SP600125, a JNK inhibitor; or 100 nM U0126, a MEK inhibitor) after 3–4 weeks of monolayer culture. SP600125 treatment significantly improved neurite outgrowth and cell viability and reduced cytotoxicity, whereas treatment with U0126 significantly enhanced neurite outgrowth and reduced cytotoxicity (n = 3). **l,** Representative neurite images of SBMA3E10 MNs before treatment (4 weeks) and after 1 week of treatment (5 weeks) with 0.0001% DMSO, 10 nM SB657510 or 1 nM GSK1562590. Neurite degeneration was observed in DMSO-treated neurons (arrows), whereas treatment with UTS2R inhibitors rescued neurite outgrowth (arrowheads). Scale bar, 100 μm. **m,** Neurite length (left), cell viability (MTS assay, middle), and cytotoxicity (LDH assay, right) of SBMA MNs treated for 1 week with 0.0001% DMSO or UTS2R inhibitors (10 nM SB657510 or 1 nM GSK1562590) after 4 weeks of monolayer culture. Treatment with UTS2R inhibitors significantly reversed neurite degeneration, increased cell viability, and reduced cytotoxicity (n = 3). The data are presented as the mean ± SEM; *, *p* < 0.05; **, *p* < 0.01; one-way repeated-measures ANOVA followed by Dunnett’s multiple comparisons test.

To assess the physiological relevance of each neuropeptide, mutant AR and each neuropeptide were knocked down using lentiviruses expressing shRNAs targeting each molecule under the control of the U6 promoter and Venus fluorescent protein under the control of the *HB9^e438^* enhancer and β*-globin* minimal promoter (Supplementary Fig. 6d and Fig. 6a). As expected, compared with shControl, knockdown of mutant AR significantly rescued neurite outgrowth by 1.36 ± 0.04-fold (Fig. 6c, n = 3, *p* < 0.01). In SBMA MNs, compared with shControl, knockdown of *CALCA*, *CGRP-1*, *UTS2*, and *UCN* significantly increased neurite outgrowth by approximately 1.27-, 1.34-, 1.43-, and 1.43-fold, respectively, at 8 weeks (Fig. 6d; n = 3; *p* < 0.05). In contrast, shCTN did not have a statistically significant effect. Consistent with the low expression of VIP in the spinal cord MNs of SBMA patients, *VIP* knockdown did not rescue neurite outgrowth (Supplementary Fig. 5a). Furthermore, knockdown of each neuropeptide did not affect neurite outgrowth in control MNs (Supplementary Fig. 6e). Together, these results suggest that CGRP-1, UTS2, and UCN, but not CTN or VIP, are functional contributors to the pathological phenotype of SBMA MNs.

### JNK and ERK1/2 constitute core signaling pathways downstream of the four neuropeptides involved in SBMA pathogenesis

We next examined whether each neuropeptide could reproduce the SBMA disease phenotype. The peptides CGRP-1, UTS2, UCN, and VIP were administered to control MNs, and their effects on neurite outgrowth were assessed. Consistent with the results of forced expression, treatment with each neuropeptide suppressed neurite outgrowth by approximately 20–35%, suggesting that these four neuropeptides contribute to SBMA pathology (Fig. 6e, f).

To investigate the downstream signaling pathways involved, control MNs were exposed to the above neuropeptides, and the phosphorylation of key signaling proteins was analyzed (Fig. 6g). Time-course immunoblotting analysis over 60 minutes revealed consistent changes in the phosphorylation of c-Jun N-terminal kinase (JNK), c-Jun, ERK1/2, and p38 MAPK across MNs derived from three independent control iPSC clones, suggesting that these signaling cascades are involved in SBMA pathogenesis (Fig. 6h, i and Supplementary Fig. 6f).

Consistent with these observations, treatment of SBMA MNs cultured for 4 weeks with an ERK1/2 pathway inhibitor (the MEK1/2 inhibitor U0126) or a JNK inhibitor (SP600125) partially rescued neurite outgrowth and reduced cytotoxicity, as determined by LDH leakage measurements (Fig. 6j, k). Compared with the control, the JNK inhibitor significantly increased cell viability (3.7 ± 1.2% increase relative to the control; n = 3; *p* < 0.05) and reduced cytotoxicity (30.1 ± 7.1% decrease relative to the control; n = 3; *p* < 0.05), whereas the MEK1/2 inhibitor increased neurite outgrowth (1.9 ± 0.2% increase relative to the control; n = 3; *p* < 0.05) and reduced cytotoxicity (32.9 ± 7.6% decrease relative to the control; n = 3; *p* < 0.05) without affecting cell viability. Neither inhibitor affected control MNs (Supplementary Fig. 6h). In agreement with previous findings in SH-SY5Y cells showing that JNK inhibition is beneficial for neurite outgrowth and cell viability, our results in SBMA patient-derived MNs support pathogenic roles for JNK and ERK1/2 as key downstream mediators of these neuropeptides^18,22^.

### UTS2 receptor antagonists partially rescue the SBMA MN phenotype

Finally, we investigated the role of UTS2, which, like CGRP-1, is highly expressed in MNs (Fig. 4g). After confirming that MNs express the UTS2 receptor (UTS2R) as well as the receptors for the other identified neuropeptides, we investigated whether pharmacological inhibition of UTS2 signaling could ameliorate the disease phenotype of SBMA MNs (Supplementary Fig. 6g). Treatment of SBMA MNs with either of the selective UTS2R antagonists SB657510 and GSK1562590 significantly rescued neurite outgrowth (by 2.9 ± 0.7% and 1.9 ± 0.2% relative to the control, respectively), modestly but significantly improved cell viability and reduced cytotoxicity without affecting control MNs (Fig. 6j, l, m and Supplementary Fig. 6i). These results suggest that UTS2 signaling contributes to SBMA pathogenesis and highlight UTS2R as a potential therapeutic target.

## Discussion

In this study, an iPSC-derived MN model successfully recapitulated early-stage SBMA pathology, including progressive neurite degeneration, upregulation of *CALCA*, and increased c-Jun phosphorylation. Moreover, many transcriptomic changes overlapped with the DEGs identified in preclinical/early-stage SBMA model mice, collectively indicating early pathological alterations. Clinically, antiandrogen therapy is most beneficial in early-stage patients, highlighting the importance of early diagnosis and intervention^12^. The identification of early-stage biomarkers in iPSC-derived MNs could facilitate early diagnosis, enable disease staging, and potentially serve as surrogate markers for assessing treatment response even before overt clinical symptoms occur.

Interestingly, although early-stage pathological features were clearly present in our iPSC-derived MNs, mutant AR aggregates were not detected. This observation is consistent with most previous reports in which mutant AR aggregates are rarely observed in iPSC-derived MNs^16,17,23,24^. To our knowledge, only a single study has revealed their detection^25^. These findings suggest that early neuronal dysfunction in the context of SBMA can occur prior to the overt aggregation of mutant AR. Given the slowly progressive nature of SBMA, disease-related stressors may be required to detect mutant AR aggregation. Future work should focus on identifying conditions that promote mutant AR oligomerization and aggregation in iPSC-derived models, thereby providing a platform to investigate disease mechanisms and therapeutic strategies at more advanced stages.

To elucidate the mechanisms underlying early-stage pathology, we examined whether the observed phenotypes depended on the AR ligand, testosterone or DHT. Although SBMA has traditionally been considered a testosterone-dependent disease, our iPSC-derived MN model exhibited early pathological phenotypes, including neurite degeneration, molecular alterations, and activation of stress-response signaling, even in the absence of DHT. These findings are consistent with those of previous studies using SBMA disease-specific iPSCs, which reported limited DHT-dependent effects, such as reduced neurite outgrowth, elevated levels of cleaved caspase-3, and altered mitochondrial membrane potential, regardless of the presence of DHT^17,23,24^. Moreover, *in vivo* studies have demonstrated abnormalities in the neuromuscular junction (NMJ) under ligand-deficient conditions in multiple SBMA female mouse models prior to disease onset and that human female carriers can present mild clinical symptoms^26–29^. Together, these findings suggest that mutant AR can exert pathogenic effects through ligand-independent mechanisms, particularly at the early stages of the disease. Given that AR functions as a nuclear receptor, ligand-independent transcriptional dysregulation by mutant AR in addition to other mechanisms, such as aggregation-driven transcriptional dysregulation and posttranslational modifications that modulate AR toxicity, may contribute to SBMA pathogenesis^5,30^.

In terms of disease progression, ER stress has emerged as a key pathogenic factor, as demonstrated in this study. ER stress has been implicated in various neurodegenerative disorders, including SBMA and several other polyglutamine diseases^14,20,31–36^. In MNs from AR-100Q mice, early increases in the levels of ER stress markers, such as BIP, ATF4, and CHOP, are accompanied by cell death^36^. Similarly, our human iPSC-derived MNs exhibited elevated expression of ATF4, CHOP, and sXBP1, reflecting activation of the PERK and IRE1 branches of the UPR, which may contribute to disease initiation or progression. Notably, ATF6 has also been reported to regulate XBP1 splicing^37,38^, indicating that crosstalk among UPR branches occurs during neurodegeneration. In SBMA, persistent ER stress driven by the misfolding and aggregation of mutant AR is thought to increase with age and due to other factors, ultimately leading to neuronal loss. In our MN model, chronic modest ER stress suppressed neurite outgrowth. These findings suggest that ER stress contributes not only to disease pathology but also to disease progression and that interventions targeting ER stress may therefore mitigate SBMA pathology and disease progression, which is consistent with reports that salubrinal treatment suppressed neuronal cell death in SBMA^36^.

Upon exploration of the molecular mechanisms underlying SBMA pathology, transcriptome analysis revealed four neuropeptides associated with early-stage disease. These neuropeptides are broadly distributed across cerebral regions, as well as in the hypothalamus and spinal cord^39–42^, and may interact with one another through shared signaling pathways^43–45^. Neuropathological examination of the lumbar spinal anterior horn confirmed the presence of CGRP, UTS2, and UCN, which were significantly more highly expressed in the MNs of SBMA patients than in those of controls. These findings are consistent with their expression levels in iPSC-derived MNs and support their pathophysiological relevance.

How might these neuropeptides, then, induce neurodegeneration? Mechanistically, they may do so by converging on stress-responsive kinase cascades. In our system, downstream activation of the JNK, c-Jun, and ERK1/2 signaling pathways, which are typically triggered by cellular stress and linked to neuronal injury, was observed. The coordinated activation of these signals via the four neuropeptides suggests partially overlapping mechanisms of action and convergence on common signaling pathways, ultimately resulting in cellular dysfunction and damage. The expression of cognate receptors for these neuropeptides in MNs supports a cell-autonomous mechanism, whereas expression in skeletal muscle indicates a noncell-autonomous component, and both modes are likely relevant to disease pathogenesis. Consistent with these observations, the expression levels of synapse-related genes were altered from early stages in SBMA MNs, as previously reported^14,46^, indicating that these neuropeptides are involved in NMJ pathogenesis.

We also evaluated the potential of these neuropeptides as biomarkers, considering their detectability *in vivo*. In our study, UCN and VIP were expressed at low levels in iPSC-derived MNs and VIP expression was similarly low in human spinal cord MNs, limiting their immediate utility as biomarkers. In contrast, CGRP and UTS2 were robustly expressed both *in vitro* and in SBMA patient spinal cord MNs. Given that they are detectable in serum, these peptides represent plausible biomarker candidates. Although they have been infrequently studied in motor neuron diseases, our findings indicate that CGRP and UTS2 are closely linked to MN pathology, with UTS2 emerging as a particularly promising therapeutic target.

*Uts2* expression was enhanced in SBMA model mice at preclinical, early, and advanced disease stages. *UTS2* encodes a neuropeptide whose mRNA is enriched in the brainstem and spinal MNs, and its receptor is closely associated with cholinergic terminals in the spinal cord, which is consistent with a role in modulating MN activity^47–49^. Notably, immunostaining showed that UTS2 colocalizes with AR^48^, and SBMA patient spinal MNs with high UTS2 expression harbored ubiquitin-positive NIs containing mutant AR, suggesting an association between mutant AR and UTS2. In our iPSC-derived MN model, pharmacological inhibition with SB657510, a selective UTS2 receptor inhibitor, attenuated neurite degeneration and modestly increased cell viability. Given that UTS2 receptors are expressed not only in MNs but also more abundantly in skeletal muscle, particularly at NMJs^47^, SB657510 may confer additional therapeutic benefits in both skeletal muscle and NMJs. Leuprorelin, the only approved pharmacological therapy for SBMA, suppresses serum testosterone levels and consequently AR function, improving SBMA symptoms but causing adverse effects associated with antiandrogen therapy, including sexual dysfunction. In contrast, SB657510 may suppress disease activity while avoiding these adverse effects. Taken together, the high expression of UTS2 in MNs and its contribution to disease pathology suggest its promise as a therapeutic target in SBMA and its potential relevance to other motor neuron diseases, such as ALS or SMA, although further investigations are needed.

## Materials and methods

### iPSC culture and differentiation

hiPSCs were maintained on mitomycin-C-treated SNL murine fibroblast feeder cells in 0.1% gelatin-coated tissue culture dishes in hESC medium and were used for motor neuron induction as described previously^13,14^. To induce differentiation, hiPSC colonies were detached using a dissociation solution (0.25% trypsin, 100 μg/ml collagenase IV (Gibco, USA), 1 mM CaCl_2_, and 20% KSR (Gibco)) and cultured in suspension in bacteriological dishes in standard hESC medium without FGF-2 after the SNL feeder cells were removed. On day 1, the medium was changed to human embryoid body (hEB) medium containing DMEM/F-12, 5% KSR, 2 mM L-glutamine, 1% NEAAs, and 0.1 mM 2-ME supplemented with 300 nM LDN-193189 (Sigma–Aldrich, USA), 3 μM SB431542 (Tocris, UK), and 3 μM CHIR99021 (FCS, USA). On day 2, the medium was changed to fresh hEB medium containing 300 nM LDN-193189, 3 μM SB431542, 3 μM CHIR99021, and 1 μM retinoic acid (RA) (Sigma–Aldrich). From day 4 to day 14, hEBs were cultured in hEB medium supplemented with 1 μM RA and 1 μM purmorphamine (Calbiochem, Germany), and the medium was changed every 2–3 days. On day 14, the hEBs were enzymatically dissociated into single cells using TrypLE Select (Thermo Fisher Scientific). The dissociated cells were plated on poly-L-ornithine (PO) and recombinant mouse laminin (Thermo Fisher Scientific) or growth factor-reduced Matrigel (Corning)-coated dishes at a density of 5×10^4^–1×10^5^ cells/cm^2^ and cultured in motor neuron medium (MNM) consisting of media containing a hormone mix (MHM)^50^ or KBM Neural Stem medium (Kohjin Bio, Japan) supplemented with B27 supplement (Thermo Fisher Scientific), 1% NEAAs, 50 nM RA, 500 nM purmorphamine, 10 μM cyclic AMP (cAMP) (Sigma–Aldrich, USA), 10 ng/mL recombinant BDNF (R&D systems), 10 ng/mL recombinant GDNF (R&D systems), and 200 ng/mL L-ascorbic acid (Sigma–Aldrich) with or without 10 ng/mL recombinant human IGF-1 (R&D systems).

### Immunocytochemical analysis

Cells were fixed in 4% paraformaldehyde for 30 min at room temperature. After blocking in blocking buffer (PBS containing 10% FBS and 0.3% Triton X-100), the cells were incubated with primary antibodies overnight at 4°C. The cells were then rinsed with PBS three times and incubated with Alexa 488-, Alexa 555-, or Alexa 647-conjugated secondary antibodies (Thermo Fisher Scientific) for 1 hour at room temperature. The nuclei were stained with 10 μg/ml Hoechst 33258 (Sigma–Aldrich). The cells were then rinsed with PBS three times, mounted on slides and examined using an IX83 (Olympus, Japan) or LSM900 (Carl Zeiss, Germany) microscope. The primary antibodies used in these analyses are listed in Supplementary Table 6.

### Western blot analysis

Western blot analysis was performed as described previously^51^. A protein sample from a total cell extract was separated by 7.5 or 10% sodium dodecyl sulfate polyacrylamide gel and transferred to a polyvinylidene fluoride (PVDF) membrane (Millipore). The blot was then probed with the primary antibodies listed in Supplementary Table 6. Signals were detected with HRP-conjugated secondary antibodies (Jackson ImmunoResearch Laboratory Inc., USA) using ECL or an ECL Prime kit (Amersham Biosciences, USA). Quantitative analysis was performed using Fusion Solo S and its software (Vilber-Lourmat, France). The amount of protein was normalized to that of α-Tubulin or β-actin. Thirty micrograms of total protein was loaded for AR detection. For the other experiments, 4 μg of total protein was loaded.

### RNA isolation and quantitative RT–PCR

RNA was isolated using an RNeasy Mini Kit (Qiagen, Germany) and then converted into cDNA using SuperScript III reverse transcriptase (Thermo Fisher Scientific) and Oligo dT primers as described previously^50,51^. To confirm the overexpression and knockdown of the targets, the CellAmp™ Direct RNA Prep Kit for RT–PCR (Takara Bio, Japan) was used according to the manufacturer’s instructions. Real-time quantitative RT–PCR was performed as previously described using SYBR Premix ExTaq II (Takara Bio) with the StepOnePlus or the QuantStudio 7 Real-Time PCR system (Thermo Fisher Scientific). The amount of cDNA was normalized to that of human-specific β*-ACTIN* mRNA. The primer sequences and PCR cycling conditions are listed in Supplementary Table 7.

### Transcriptome analysis

The experimental procedure for the cDNA microarray analysis was based on the manufacturer’s protocol (Agilent Technologies, USA). In brief, cDNA synthesis and cRNA labeling with cyanine-3 dye were conducted using an Agilent Low Input Quick Amp Labeling Kit (Agilent Technologies). Cyanine 3-labeled cRNA was purified, fragmented, and hybridized on a Human Gene Expression 4 × 44K v2 Microarray Chip (G2519F#026652) containing 27,958 Entrez Gene RNAs using a Gene Expression Hybridization kit (Agilent Technologies). Gene expression in a set of SBMA3E10 and TIGE9 MNs was analyzed at 4 weeks. The experiment was repeated twice (2 sets), yielding 4 samples. The expression data were log-transformed and normalized with a 75th percentile shift using GeneSpring (Agilent Technologies). The criterion used to detect the differences in gene expression was a 1.5-fold change from that of 26,308 genes (28,961 probes). The raw and normalized microarray data have been submitted to the NCBI GEO database (accession no. GSE310066). We also used mouse microarray data reported previously (accession no. GSE39865). Briefly, total spinal cords of wild-type, AR-24Q and AR-97Q male mice aged 7–9 weeks (before disease onset), 10–12 weeks (early disease stage) and 13–15 weeks (advanced disease stage), for a total of 21 samples, were examined by microarray analysis using an Affymetrix GeneChip system (Mouse Genome 430 2.0 Array (Affymetrix, Thermo Fisher Scientific)). The raw data were filtered with a lower cutoff of 20% and an upper cutoff of 100%, and at least 1 out of 21 samples had values within the cutoff. We normalized the raw data with a 75^th^ percentile shift and identified significantly different probe sets (*p* < 0.05) by performing a moderated *t* test with the Westfall–Young procedure for the 18,678 genes (39,403 probes). The expression data were grouped using a hierarchical clustering algorithm with the Gene Cluster 3.0 program^52^. A heatmap was generated using the Java TreeView program^53^.

### ELISA

The supernatants from the iPSC-derived MN cultures were collected, and we used CGRP EIA kits, CGRP Fluorescent EIA kits, Urotensin II Fluorescent EIA kits, Urocortin Fluorescent EIA kits, and VIP Fluorescent EIA kits (Phoenix Pharmaceuticals, USA) according to the manufacturer’s instructions for analysis.

### Preparation of lentiviruses

Motor neuron reporter lentivirus (*HB9^e438^::Venus*) was prepared as described previously ^13^. In brief, Lenti-X 293T cells (Takara Bio) were transfected with pSIN2-HB9^e438^-βglo-Venus together with packaging plasmids (pCMV-VSV-G-RSV-Rev and pCAG-HIV-gp, kindly provided by late Dr. Hiroyuki Miyoshi). Cells were cultured in the presence of 10 μM forskolin (Cayman), and the conditioned medium containing viral particles was harvested and concentrated by ultracentrifugation at 25,000 rpm for 90 minutes at 4°C. To generate the overexpression vectors of the neuropeptides, cDNAs encoding the four neuropeptides were amplified by PCR using the primers listed in Supplementary Table 8, cloned and inserted into a pENTR vector (Thermo Fisher Scientific). The inserts were then transferred into a lentiviral destination vector (pSIN2-HB9^e438^-βglo-RfC.1-IRES2-Venus) using the Gateway system (Thermo Fisher Scientific) to generate pSIN2-HB9^e438^-βglo-GOI-IRES2-Venus. Lentiviruses expressing neuropeptides (*HB9^e438^::GOI-IRES-Venus*) were produced using the same procedure as that used for the reporter virus.

For the preparation of shRNA-expressing vectors, target sequences were designed using siDirect (https://sidirect2.rnai.jp/v2.0/), BLOCK-iT RNAi Designer (https://rnaidesigner.thermofisher.com/rnaiexpress/), or the shRNA Target Design tool of VectorBuilder (https://en.vectorbuilder.com/tool/shrna-target-design.html). The designed oligonucleotides (listed in Supplementary Table 8) were annealed, cloned and inserted into pENTR-U6-shRNA, and the resulting constructs were transferred into the lentiviral destination vector (pCS-RfA-*HB9^e438^*-βglo-Venus lentivirus) using the Gateway system to generate pCS-U6-shRNA-HB9^e438^-βglo-Venus plasmids. Lentiviruses expressing shRNA were produced as described above, except that the conditioned medium containing viral particles was directly used for transduction without ultracentrifugation.

For lentiviral titration, viral RNA was extracted using NucleoSpin RNA Virus (Macherey-Nagel, Germany), and RNA genome copy numbers were quantified with a Lenti-X qRT–PCR Titration Kit (Takara Bio) according to the manufacturer’s instructions.

### Neurite length assay

iPSC-derived MNs were plated on Matrigel-coated 96-well imaging plates (655090, Greiner, Germany) at a density of 1.5 × 10^4^ cells/well in the presence of CultureOne Supplement (Thermo Fisher Scientific) and cultured for 4–8 weeks. Lentiviral transduction was performed between days 2 and 4 of monolayer MN differentiation. A time course analysis of neurite length was performed over 8 weeks at intervals of 3–7 days. Images of 4 × 4 fields were acquired from each well using a 10× objective lens on an IN Cell Analyzer 6000 (Cytiva, USA). Neurite length was quantified using IN Cell Developer Toolbox 1.9.2 (Cytiva), and the values at each time point were normalized to that measured on the first day. The peptides, receptor inhibitors, and other compounds used in this study are listed in Supplementary Table 9.

### Cell viability and cytotoxicity assays

Cell viability and cytotoxicity were assessed via the CellTiter 96 AQueous One Solution Cell Proliferation Assay (MTS assay; Promega, USA) and the Cytotoxicity LDH Assay Kit-WST (Dojindo, Japan), respectively, according to the manufacturers’ instructions. For the cell viability assay, cells cultured in 96-well plates were incubated with MTS reagent for 3 hours, and the absorbance was measured at 490 nm with background subtraction at 655 nm using an iMark microplate reader (Bio-Rad, USA). For the LDH cytotoxicity assay, 50–100 μL of culture medium was incubated with the substrate for 30 minutes, after which the absorbance was measured at 490 nm.

### Pathological assessment of autopsied SBMA patients

Human autopsy spinal cord samples were obtained with written informed consent from the patients’ families and used in accordance with protocols approved by the Ethics Committee of Aichi Medical University. Spinal cords were fixed in 20% formalin for at least one month and subsequently embedded in paraffin. Axial sections from all the patients were prepared and subjected to immunohistochemical analysis. The primary antibodies used are listed in Supplementary Table 6. Antigen retrieval was performed in citrate buffer (pH 6.0) at 98°C for 25 minutes. Secondary immunolabeling was performed using a standard avidin-biotin method. Double immunofluorescence staining was performed using an anti-ubiquitin antibody together with anti-CGRP, anti-UTS2, or anti-UCN antibodies. Alexa 488- or 568-conjugated anti-mouse or anti-rabbit IgG (Thermo Fisher Scientific) was used for secondary labeling. The immunofluorescent samples were observed using a laser scanning confocal microscope (FV-3000; Olympus, Tokyo, Japan) with identical settings for each combination of primary antibodies.

The fluorescence intensity of the MNs in the anterior horn of the spinal cord was quantified from photomicrographs obtained by confocal microscopy. Sections (10 μm thick) were prepared for analysis. Four visual fields at 400× magnification per lumbar segment, each at least 200 μm apart, were randomly selected from two lumbar cord segments. The region of interest (ROI) of each neuron was defined as the cytoplasmic area minus the nuclear and lipofuscin areas. The output was the mean signal intensity within the ROI. Measurements were performed at the focal plane corresponding to the largest nuclear diameter, and neurons lacking nuclei within that plane were excluded from the analysis.

### Analysis of the number of CAG repeats in paraffin-embedded tissue

Genomic DNA was extracted from paraffin-embedded tissue using the NucleoSpin DNA FFPE XS kit (Macherey-Nagel). The CAG repeat region of the *AR* gene was amplified by PCR, followed by direct sequencing to determine the number of CAG repeats. The primer sequences and PCR cycling conditions are listed in Supplementary Table 7.

### Statistical analysis

The data are expressed as the mean ± standard error of the mean (SEM). For the qRT–PCR data, statistical significance was assessed using either Student’s *t* test or Welch’s *t* test. The time course neurite length analysis data were analyzed using repeated measures one-way or two-way analysis of variance (ANOVA) followed by the Tukey–Kramer or Dunnett multiple comparison test. For quantification of the immunohistochemistry results, the Kruskal–Wallis test was used.

## Supporting information

Supplementary Table2 and 3_Up and Down

## Ethics approval and consent to participate

All experimental procedures for the establishment and use of human iPSCs were approved by the Ethics Committee of Keio University School of Medicine (approval number 20080016) and the Ethics Committee of Aichi Medical University School of Medicine (approval numbers 2023-H006 and 2020-213). Autopsies were performed after written informed consent from family members and approval from the Ethics Committee of Aichi Medical University School of Medicine (approval number 2019M019) were obtained in accordance with institutional ethical guidelines.

## Availability of the data and materials

The datasets supporting the conclusions of this article are available in the following repositories: the expression data have been deposited in the GEO [http://www.ncbi.nlm.nih.gov/geo/]

## Acknowledgments

We are grateful to Dr. S. Yamanaka (Kyoto University) for supplying human iPSCs (201B7); to the late Dr. H. Miyoshi (RIKEN BRC and Keio University) for providing the lentiviral vectors; to the Aichi Medical University Institute of Comprehensive Medical Research Division of Advanced Research Promotion for assistance with the experiments; to Ms. M. Ishihara, Ms. Y. Kitashiba, Ms. S. Tokuda, Ms. S. Kumagai, Ms. E. Kojima, Ms. S. Yokoi, and Ms. Y. King for technical and administrative support; and to all members of Dr. Okada’s laboratory for encouragement and support.

## Author contributions

KO and YO conceived and designed the study. KO, DS, MIR and YO performed the experiments and analyzed the data. KO, AO and YH performed the microarray analysis. YR, MY and YI performed the histological analysis of the autopsied samples. SY, MK and GS recruited SBMA patients. RO and SY provided technical assistance. KO and YO wrote the manuscript. MD, GS, MK, and HO provided scientific discussions and critically reviewed the manuscript. All the authors read and approved the final manuscript.

