## Supplementary_information for "Early pathogenesis of spinal and bulbar muscular atrophy uncovered by human iPSC-derived motor neurons highlights pathogenic neuropeptides as therapeutic targets"

**Supplementary Tables**

Supplementary Table 1. Genes (probes) extracted from the mouse microarray.
Supplementary Table 2. (in Excel file) List of upregulated genes in human SBMA iPSC-derived MNs.
Supplementary Table 3. (in Excel file) List of downregulated genes in human SBMA iPSC-derived MNs.
Supplementary Table 4. List of upregulated and downregulated genes in both human SBMA iPSC-derived MNs and AR97Q mice.
Supplementary Table 5. Demographic information of the autopsied patients.
Supplementary Table 6. Antibodies used in this study.
Supplementary Table 7. Primers used for RT–PCR and CAG repeat sizing.
Supplementary Table 8. Primers used for cloning and shRNA target sequences.
Supplementary Table 9. Peptides and compounds used in this study.

**Supplementary Figures**

Supplementary Figure 1. Differentiation and maturation of control and SBMA iPSCs into MNs.
Supplementary Figure 2. Neuronal cell death and cytotoxicity.
Supplementary Figure 3. PolyQ expression under ER stress.
Supplementary Figure 4. Human and mouse transcriptome analysis results.
Supplementary Figure 5. VIP is rarely detected in spinal MNs from control or SBMA patients.
Supplementary Figure 6. Validation of lentiviral vectors and the effects of the neuropeptides on control MNs.

**Supplementary Table 1.**
**Genes (probes) extracted from the mouse microarray.**

| | Up | Down | Total |
|---|---|---|---|
| Before onset (preclinical) | 168 (458) | 184 (331) | 316 (789) |
| Early | 78 (149) | 60 (118) | 138 (267) |
| Advanced | 48 (131) | 116 (224) | 164 (355) |
| Total | 294 (738) | 360 (673) | 618 (1411) |

**Supplementary Table 4.**
**List of upregulated genes in both human SBMA iPSC-derived MNs and AR97Q mice.**

| | SBMA/control_FC | | Mouse AR97Q/AR24Q_FC | | |
|---|---|---|---|---|---|
| Gene symbol | Experiment 1 | Experiment 2 | Preclinical | Early | Advanced |
| *Acacb* | 2.88 | 1.72 | 1.32* | 0.97 | 0.88 |
| *Agmo* | 4.58 | 4.09 | 0.77 | 1.27* | 1.20 |
| *Atf7* | 2.12 | 2.02 | 0.98 | 1.29* | 1.01 |
| *C1ql3* | 1.80 | 2.31 | 1.16 | 1.00 | 1.28* |
| *C1ql3* | 1.80 | 2.31 | 1.26 | 0.93 | 1.29* |
| *Calca* | 2.03 | 1.05 | 1.51 | 2.34* | 3.45* |
| *Ccdc114* | 1.12 | 3.82 | 0.95 | 1.24* | 0.90 |
| *Crem* | 1.58 | 1.27 | 0.93 | 1.00 | 1.50* |
| *Ctsf* | 1.23 | 1.75 | 1.27* | 1.00 | 1.11 |
| *Epb41l3* | 1.72 | 1.72 | 1.28* | 1.02 | 1.11 |
| *Fam120c* | 1.64 | 1.21 | 1.45* | 0.96 | 1.06 |
| *Fam227b* | 1.60 | 1.34 | 0.83 | 1.66* | 0.78 |
| *Gas6* | 1.49 | 1.83 | 1.26* | 0.97 | 1.05 |
| *Ltbp3* | 1.20 | 2.09 | 1.42* | 1.06 | 1.05 |
| *Mall* | 1.84 | 1.96 | 1.23 | 1.34* | 0.88 |
| *Max* | 1.51 | 1.31 | 1.06 | 1.47* | 1.05 |
| *Mier1* | 1.61 | 1.01 | 1.35* | 1.03 | 1.19 |
| *Nadk2* | 1.58 | 1.19 | 0.89 | 1.26* | 1.13 |
| *Nfib* | 1.79 | 1.14 | 1.25* | 1.06 | 0.97 |
| *Pag1* | 1.50 | 1.31 | 0.97 | 1.24* | 0.84 |
| *Pappa* | 1.88 | 1.78 | 1.47* | 1.06 | 1.43* |
| *Pappa* | 1.88 | 1.78 | 1.41 | 0.97 | 1.49* |
| *Pde1c* | 2.23 | 1.27 | 1.32* | 1.08 | 1.00 |
| *Pecam1* | 1.59 | 1.17 | 1.41* | 0.94 | 1.05 |
| *Pramef12* | 2.10 | 1.32 | 1.07 | 1.28* | 1.08 |
| *Rapgef3* | 1.61 | 1.12 | 1.75* | 0.84 | 1.05 |
| *Ret* | 2.56 | 1.79 | 1.26* | 1.01 | 1.09 |
| *Scrg1* | 1.96 | 1.54 | 1.20 | 1.06 | 1.44* |
| *Sdc4* | 1.50 | 2.28 | 1.30* | 1.11 | 1.02 |
| *Selenbp1* | 2.95 | 2.68 | 1.35* | 1.11 | 1.11 |
| *Selenbp1* | 2.95 | 2.68 | 1.37* | 1.06 | 1.16 |

| | | | | | |
|---|---|---|---|---|---|
| *Slc4a4* | 1.82 | 1.38 | 1.59* | 0.84 | 1.16 |
| *Slc4a4* | 1.82 | 1.38 | 1.36* | 0.87 | 1.00 |
| *Slc5a7* | 1.14 | 2.01 | 1.79* | 0.94 | 1.53* |
| *Spink4* | 1.29 | 2.03 | 1.00 | 1.25* | 0.89 |
| *Spon1* | 4.46 | 1.66 | 1.45* | 0.97 | 1.18 |
| *Tapbp* | 1.30 | 1.59 | 1.31* | 1.15 | 1.16 |
| *Tbl1x* | 1.25 | 1.57 | 1.47* | 0.92 | 0.98 |
| *Timp3* | 2.20 | 1.21 | 1.40* | 1.03 | 0.95 |
| *Twist2* | 1.79 | 1.08 | 0.82 | 1.24* | 0.93 |
| *Ubash3b* | 1.72 | 1.00 | 1.35* | 1.03 | 0.98 |
| *Ubxn11* | 1.39 | 1.82 | 1.07 | 1.27* | 0.99 |
| *Ucn* | 2.46 | 1.48 | 2.02 | 2.19* | 3.29* |
| *Urb1* | 1.48 | 1.68 | 1.11 | 1.01 | 1.37* |
| *Uts2* | 2.24 | 2.83 | 3.52* | 2.58* | 4.70* |
| *Vip* | 1.60 | 2.14 | 1.61 | 1.59* | 2.40* |

**List of downregulated genes in both human SBMA iPSC-derived MNs and AR97Q mice.**

| | SBMA/control_FC | | Mouse AR97Q/AR24Q_FC | | |
|---|---|---|---|---|---|
| Gene symbol | Experiment 1 | Experiment 2 | Preclinical | Early | Advanced |
| *Acbd5* | 0.57 | 0.93 | 0.75* | 1.06 | 0.97 |
| *Ackr3* | 0.62 | 0.99 | 0.62* | 0.86 | 0.67 |
| *Adam33* | 0.45 | 1.00 | 0.95 | 0.55* | 1.16 |
| *Adamtsl2* | 0.96 | 0.61 | 1.04 | 1.01 | 0.79* |
| *Ahsp* | 0.61 | 0.98 | 0.50* | 0.60* | 0.65 |
| *Cadm2* | 0.79 | 0.48 | 0.75* | 1.09 | 1.12 |
| *Cadm2* | 0.79 | 0.48 | 0.77* | 1.15 | 0.97 |
| *Chn2* | 0.94 | 0.36 | 0.74 | 0.74* | 1.06 |
| *Chrdl1* | 0.68 | 0.48 | 0.72* | 1.02 | 0.83 |
| *Col13a1* | 0.68 | 0.56 | 0.96 | 1.04 | 0.67* |
| *Col15a1* | 0.74 | 0.47 | 0.68* | 0.82 | 0.96 |
| *Col1a2* | 0.18 | 0.11 | 0.37* | 0.78 | 0.64 |
| *Col3a1* | 0.09 | 0.21 | 0.32* | 0.70 | 0.52 |
| *Col3a1* | 0.09 | 0.21 | 0.66* | 0.79 | 0.77* |
| *Foxd3* | 0.71 | 0.29 | 0.96 | 1.14 | 0.79* |
| *Hpgd* | 0.26 | 0.66 | 0.75* | 0.88 | 0.92 |
| *Iglon5* | 0.96 | 0.62 | 0.63* | 1.30 | 0.82 |
| *Lamb1* | 0.76 | 0.04 | 0.61* | 0.82 | 0.83 |
| *Lamb1* | 0.76 | 0.04 | 0.66* | 0.93 | 0.82 |
| *Ncan* | 0.99 | 0.55 | 0.74* | 0.87 | 0.86 |
| *Ntsr2* | 0.31 | 0.99 | 0.93 | 0.88 | 0.81* |
| *Plxdc1* | 0.23 | 0.33 | 0.98 | 0.74* | 0.82 |
| *Prg2* | 0.38 | 0.55 | 0.74* | 0.61* | 1.01 |
| *Sh3rf1* | 0.83 | 0.55 | 0.98 | 1.14 | 0.80* |
| *Slc15a4* | 0.84 | 0.34 | 0.94 | 0.79* | 1.00 |
| *Sox18* | 0.59 | 0.79 | 1.01 | 0.74* | 0.83 |

| | | | | | |
|---|---|---|---|---|---|
| *Srcin1* | 0.53 | 0.90 | 0.76 | 0.89 | 0.77* |
| *Them5* | 0.63 | 0.74 | 0.70* | 1.12 | 0.78 |
| *Trhr* | 0.92 | 0.63 | 0.88 | 0.78* | 0.88 |
| *Trhr* | 0.92 | 0.63 | 0.98 | 0.78* | 0.94 |

*FC,* fold change; * corrected $p < 0.05$

**Supplementary Table 5.**
**Demographic information of the autopsied patients.**

| Patient | Age at death | Sex | Duration (years) | Number of CAG repeats |
|---|---|---|---|---|
| SBMA case 1 | 63 | m | 43 | 45–47 CAGs* |
| SBMA case 2 | 74 | m | 24 | 47 CAGs* |
| SBMA case 3 | 65 | m | 35 | 47 CAGs† |
| Control 1 | 74 | m | - | 24–25 CAGs* |
| Control 2 | 59 | m | - | 9 CAGs* |

* from FFPE samples; † from clinical data

**Supplementary Table 6.**
**Antibodies used in this study.**

| Antibody | Clone | Host | Isotype | Dilution for ICC | Dilution for IHC | Catalog number | Source |
|---|---|---|---|---|---|---|---|
| AR | | Rb | IgG | 1:500 | | PA5-16750 | Thermo Fisher Scientific |
| βIII-tubulin | | Rb | IgG | 1:2000 | | PRB-435P | COVANCE (BioLegend) |
| βIII-tubulin | Tuj1 | Ms | IgG2a | 1:2000 | | MMS-435P | COVANCE (BioLegend) |
| CGRP | 4901 | Ms | IgG1 | 1:200 | 1:500 | C7113 | Sigma–Aldrich |
| ChAT | | Gt | IgG | 1:200 | | AB144 | Millipore |
| Cleaved Caspase-3 | | Rb | IgG | 1:5000 | | 9661 | Cell Signaling Technology |
| GFP | | Rb | IgG | 1:500 | | 598 | MBL |
| GFP | | Gt | IgG | 1:1000 | | 600-101-215 | Rockland |
| HB9 | 81.5C10 | Ms | IgG1 | 1:1000 | | 81.5C10 | Developmental Studies Hybridoma Bank [DSHB] |
| Isl-1 | 39.4D5 | Ms | IgG2b | 1:400 | | 39.4D5 | Developmental Studies Hybridoma Bank [DSHB] |
| Nanog | | Rb | IgG | 1:100 | | RCAB003P | ReproCELL |
| Oct-4 | C-10 | Ms | IgG | 1:100 | | sc-5279 | Santa Cruz Biotechnology |
| Ubiquitin | Ubi-1 | Ms | IgG1 | 1:200 | 1:200 | MAB1510 | Millipore |
| Ubiquitin | | Rb | IgG | | 1:200 | Z0458 | DAKO |
| Urocortin (UCN) | | Rb | IgG | 1:100 | 1:500 | SAB4300739 | Sigma–Aldrich |
| UTS2 | | Rb | IgG | 1:200 | 1:500 | HPA017000 | Sigma–Aldrich |
| VIP | 02 | Ms | IgG1 | 1:200 | 1:500 | ab30680 | abcam |

**Antibodies used for Western blot analysis.**

| Antibody | Clone | Host | Isotype | Dilution | Catalog number | Source |
|---|---|---|---|---|---|---|
| Akt | C67E7 | Rb | IgG | 1:1000 | 4691 | Cell Signaling Technology |

| | | | | | | |
|---|---|---|---|---|---|---|
| AR | | Rb | IgG | 1:1000 | PA5-16750 | Thermo Fisher Scientific |
| ATF6 | 70B1413.1 | Ms | IgG1 | 1:1000 | NBP1-40256 | NOVUS Biologicals |
| α-tubulin | DM1A | Ms | IgG1 | 1:2000 | T9026 | Sigma–Aldrich |
| β-actin | AC-15 | Ms | IgG1 | 1:2000 | A1978 | Sigma–Aldrich |
| c-Jun | 60A8 | Rb | IgG | 1:1000 | 9165 | Cell Signaling Technology |
| CREB | 48H2 | Rb | IgG | 1:1000 | 9197 | Cell Signaling Technology |
| eIF2a | D7D3 | Rb | IgG | 1:1000 | 5324 | Cell Signaling Technology |
| IRE1a | 14C10 | Rb | IgG | 1:1000 | 3294 | Cell Signaling Technology |
| NF-kB p65 | D14E12 | Rb | IgG | 1:1000 | 8242 | Cell Signaling Technology |
| p38 MAPK | D13E1 | Rb | IgG | 1:1000 | 8690 | Cell Signaling Technology |
| p44/22 MAPK (Erk1/2) | 137F5 | Rb | IgG | 1:1000 | 4695 | Cell Signaling Technology |
| p62 | 5F2 | Ms | IgG1 | 1:1000 | M162-3 | MBL |
| PDI | C81H6 | Rb | IgG | 1:1000 | 3501 | Cell Signaling Technology |
| PERK | D11A8 | Rb | IgG | 1:1000 | 5683 | Cell Signaling Technology |
| Phospho-Akt (Ser473) | D9E | Rb | IgG | 1:1000 | 4060 | Cell Signaling Technology |
| Phospho-c-Jun (Ser63) | 54B3 | Rb | IgG | 1:1000 | 2361 | Cell Signaling Technology |
| Phospho-CREB (Ser133) | 87G3 | Rb | IgG | 1:1000 | 9198 | Cell Signaling Technology |
| Phospho-eIF2a (Ser51) | 119A11 | Rb | IgG | 1:1000 | 3597 | Cell Signaling Technology |
| Phospho-IRE1a (p-Ser724] | | Rb | IgG | 1:1000 | NB100-2323 | NOVUS Biologicals |
| Phospho-NF-kB p65 (Ser536) | 93H1 | Rb | IgG | 1:1000 | 3033 | Cell Signaling Technology |
| Phospho-PERK (Thr980) | 16F8 | Rb | IgG | 1:1000 | 3179 | Cell Signaling Technology |
| Phospho-SAPK/JNK (Thr183/Tyr185) | | Rb | IgG | 1:1000 | 9251 | Cell Signaling Technology |
| Phospho-Smad2 (Ser465/467)/Smad3 (Ser423/425) | D27F4 | Rb | IgG | 1:1000 | 8828 | Cell Signaling Technology |
| Phospho-Src (Tyr416) | D49G4 | Rb | IgG | 1:1000 | 6943 | Cell Signaling Technology |
| Phospo-p38 MAPK (Thr180/Tyr182) | D3F9 | Rb | IgG | 1:1000 | 4511 | Cell Signaling Technology |
| Phossho-p44/22 MAPK (Erk1/2) (Thr202/Tyr204) | D13.14.4E | Rb | IgG | 1:1000 | 4370 | Cell Signaling Technology |
| Polyglutamine | 5TF1-1C2 | Ms | IgG1k | 1:1000 | MAB1574 | Sigma–Aldrich |
| SAPK/JNK | | Rb | IgG | 1:2000 | 9252 | Cell Signaling Technology |
| Smad2/3 | | Rb | IgG | 1:1000 | 5678 | Cell Signaling Technology |
| Src | | Rb | IgG | 1:1000 | 2108 | Cell Signaling Technology |

*Gt* goat, *Ms* mouse, *Rb* rabbit

**Supplementary Table 7.**
**Primers used for RT–PCR.**

| Gene | Temp. (°C) | Sense | 5' to 3' | Antisense | 5' to 3' |
|---|---|---|---|---|---|
| *AR* | 62 | hAR-S3285 | CCTGGCTTCCGCAACTTACAC | hAR-AS3452 | GGACTTGTGCATGCGGTACTCA |
| *ATF4* | 62 | hATF4_s1031 | GTTCTCCAGCGACAAGGCTA | hATF4_as1197 | TCTGGCATGGTTTCCAGGTC |
| *β-ACTIN* | 62 | hb-ACTIN-S1062 | GATCAAGATCATTGCTCCTCCT | hb-ACTIN-AS1241 | GGGTGTAACGCAACTAAGTCA |
| *BIP* | 62 | hBIP-s2134 | TGTTCAACCAATTATCAGCAAACTC | hBIP-as2206 | TTCTGCTGTATCCTCTTCACCAGT |
| *CALCA* | 62 | hCALCA-s69 | AGGTGTCATGGGCTTCCAAAAG | hCALCA-as259 | TCATCTGCACATAGTCCTGCAC |
| *CHAT* | 62 | hChAT-S1360 | GGAGGCGTGGAGCTCAGCGACACC | hChAT-AS1592 | CGGGGAGCTCGCTGACGGAGTCTG |
| *CHOP* | 62 | hCHOP-s481 | AGAACCAGGAAACGGAAACAGA | hCHOP-as547 | TCTCCTTCATGCGCTGCTTT |
| *DYNACTIN1* | 62 | hDYNACTIN-1370S | CTTGGAAGCGATGAATGAGA | hDYNACTIN-1519AS | TAGTCTGCAACCGTCTCCTG |
| *EDEM* | 56 | hEDEM_s1139 | TTTGGAACCTCCGGAGCAAT | hEDEM_as1226 | AGGCCACTCTGCTTTCCAAC |
| *HB9* | 62 | hHB9-S951 | GTCCACCGCGGGCATGATCC | hHB9-AS1162 | TCTTCACCTGGGTCTCGGTGAGC |
| *HRD1* | 62 | hHRD1_s961 | TGTCTGCATCATCTGCCGAG | hHRD1_as1099 | ACGAAGGACATCCATACGGC |
| *ISL1* | 60 | hISL1-S1294 | AGCAGCCCAATGACAAAACT | hISL1-AS1494 | CTGAAAAATTGACCAGTTGCTG |
| *sXBP1* | 62 | hsXBP-1-s533 | CTGAGTCCGAATCAGGTGCAG | hXBP-1-as591 | ATCCATGGGGAGATGTTCTGG |
| *TβRII* | 62 | hTGFBRII-2897S | TACCTTCATGGGTTGCAGAA | hTGFBRII-3030AS | AAGGCTGGGAGCAGAGAATA |
| *tXBP1* | 62 | htXBP-1-s521 | TGGCCGGGTCTGCTGAGTCCG | hXBP-1-as591 | ATCCATGGGGAGATGTTCTGG |
| *UCN* | 68 | hUCN-s543 | CGGGACAACCCTTCTCTGTC | hUCN-as669 | TGCCCACCGAGTCGAATATG |
| *UTS2* | 68 | hUTS2_s269 | TACTGCCAGAGATGCTGGGT | hUTS2_as456 | ATCAGGAGTCTCACGTTTCTTGT |
| *VIP* | 66 | hVIP-s232 | TCTCACAGACTTCGGCATGG | hVIP-as471 | TGGCAGAAAGTTGACCCAAG |

**Primers used for CAG repeat sizing.**

| Gene | Temp. | Sense | 5' to 3' | Antisense | 5' to 3' |
|---|---|---|---|---|---|
| *AR* | 68 | AR_exon1_CAG_s1182 | TCC AGA ATC TGT TCC AGA GCG TGC | AR_exon1_CAG_as1680 | TGG CCT CGC TCA GGA TGT CTT TAA G |
| | 68 | | | AR_exon1_CAG_as_1432 | CCT GTG GGG CCT CTA CGA TG |

**Supplementary Table 8.**

| | Primer used for cloning (5' to 3') |
|---|---|
| hCalcitonin-CGPR1_fw | AGAGGTGTCATGGGCTTCCA |

| | |
|---|---|
| hCalcitonin_rv | TTAGTTGGCATTCTGGGGCA |
| hCGRP1_rv | TCAGGCTTGAAGGTCCCTGC |
| hUTS2_fw | GTGGCGATCATGTATAAGCTGG |
| hUTS2_rv | TCAGACACAGTATTTCCAGA |
| hUCN_fw | GGCGGCACCATGAGGCAGGC |
| hUCN_rv | TCACTTGCCCACCGAGTCGA |
| hVIP_fw | GGCACAGAAATGGACACCAG |
| hVIP_rv | TCATTTTTCTAACTCTTCTGGA |

| | shRNA target sequence (5' to 3') |
|---|---|
| shControl2 | CGTAATACGTTCGAGGAAT |
| shControl3 | ACTTGATCGTGTATGAGGT |
| shControl4 | CCTAAGGTTAAGTCGCCCTCG |
| shAR | GAGCGTGGACTTTCCGGAAAT |
| shCALCA | GAGGTGTCATGGGCTTCCAAA |
| shCGRP1 | GAAAGGAACTATTGCTAAATG |
| shCalcitonin | ACTTGCATGCTGGGCACATAC |
| shUTS2 | GAAATTTGAGAAAGTTTCAGG |
| shUCN | GAGCAGAACCGCATCATATTC |
| shVIP | CCAGTCAAACGTCACTCAGAT |

**Supplementary Table 9.**
**Peptides and compounds used in this study.**

| Product | | Catalog number | Source |
|---|---|---|---|
| CGRP-1 | α-Calcitonin Gene Related Peptide | 4232 | Peptide Institute, Inc. |
| GSK1562590 | Selective urotensin II receptor antagonist | B7752 | APEXBIO |
| SB657510 | Selective urotensin II receptor antagonist | S7326 | Sigma–Aldrich |
| SP-600125 | Broad spectrum JNK inhibitor | S5567 | Sigma–Aldrich |
| Thapsigargin | ER stress inducer | 209-17281 | FUJIFILM Wako |
| Tunicamycin | ER stress inducer | 202-08241 | FUJIFILM Wako |
| U0126 | MEK1/2, ERK1/2 inhibitor | 211-01051 | FUJIFILM Wako |
| Urocortin (UCN) | Ligand for type 1/type 2 CRF receptors | 4328 | Peptide Institute, Inc. |
| Urotensin II (UTS2) | Potent vasoconstrictor | 4365 | Peptide Institute, Inc. |
| VIP | Vasoactive Intestinal Peptide | 4110 | Peptide Institute, Inc. |

**Supplementary Figure 1. Differentiation and maturation of control and SBMA iPSCs into MNs.**
**a,** Immunocytochemical analysis of HB9, ISL-1, βIII-Tubulin, and ChAT expression in MNs derived from control iPSCs (EKN3 and YFE16) and SBMA iPSCs (SBMA2E16 and SBMA4E5) after 1 week of monolayer differentiation.
**b,** Immunocytochemical analysis of AR expression in MNs derived from control iPSCs (EKN3 and YFE16) and SBMA iPSCs (SBMA2E16 and SBMA4E5) after 4 weeks of monolayer differentiation.
**c, d,** Western blot analysis of ChAT protein expression after 4 weeks of monolayer differentiation (**c**) and quantification of the results (**d**).
**e,** Western blot analysis of AR protein expression (related to Fig. 1l). Upper panel: Short-term exposure; lower panel: long-term exposure.
Data are presented as the mean ± SEM; *, $p < 0.05$; **, $p < 0.01$; Student's *t* test.

**Supplementary Figure 2. Neuronal cell death and cytotoxicity.**
**a,** Immunocytochemical analysis of cleaved caspase-3 expression after 2 weeks of monolayer differentiation in MNs derived from control iPSCs (EKN3 and YFE16) and SBMA iPSC clones (SBMA2E16 and SBMA4E5). Scale bar, 100 μm.
**b,** LDH assay cytotoxicity analysis using culture supernatants from MNs after 4 weeks of monolayer differentiation. No significant differences were observed between the control MNs and SBMA MNs.

**Supplementary Figure 3. PolyQ expression under ER stress.**
**a,** Western blot analysis of AR and polyglutamine (polyQ) expression in control and SBMA MNs after 4 weeks of differentiation.
**b,** Quantification of the polyQ signals shown in (**a**). The overlap of the PolyQ and AR signals markedly increased in tunicamycin (TM)-treated SBMA MNs.
**c,** Quantitative RT–PCR analysis of UPR-related genes after 4 weeks of MN differentiation (n = 3).
The data are presented as the mean ± SEM.

**Supplementary Figure 4. Human and mouse transcriptome analysis results.**
**a,** Strategy for the human and mouse transcriptome analyses. For the human dataset, genes showing more than a 1.5-fold change in expression in either or both microarray comparisons (control: TIGE9 vs. SBMA: 3E10) were included. For the mouse dataset, genes whose expression significantly changed (on the basis of a corrected $p < 0.05$) were included.
**b,** Venn diagram showing genes commonly downregulated in human SBMA patients and SBMA model mice at the preclinical, early, and advanced stages.
**c,** Hierarchical clustering analysis of the downregulated genes. The rows in the heatmap represent the selected genes, and the columns represent the mouse groups. P, preclinical; E, early; A, advanced; WT, wild type; 24Q, AR24Q; and 97Q, AR97Q.

**Supplementary Figure 5. VIP is rarely detected in spinal MNs from control or SBMA patients.**
**a,** Immunohistochemical analysis of the anterior horn of the lumbar spinal cord from SBMA patients and healthy controls. VIP staining was weak or absent in the spinal MNs from both groups. Scale bar, 100 μm.

**Supplementary Figure 6. Validation of lentiviral vectors and the effects of the neuropeptides on control MNs.**
**a,** Quantitative RT–PCR analysis of *CALCA* (CGRP-1)*, UCN, UTS2* and *VIP* expression in control MNs (EKN3, TIGE9, and YFE16) transduced with lentiviruses expressing the corresponding genes.
**b,** ELISA of CGRP-1, UTS2, UCN and VIP expression in control MNs transduced with the corresponding lentiviruses.
**c,** LDH assay of control MNs overexpressing CGRP-1, UTS2, UCN, or VIP after 4 weeks of monolayer culture.
**d,** Quantitative RT–PCR analysis of the knockdown efficiency in SBMA3E10 MNs transduced with lentivirus expressing shRNAs targeting *AR, CGRP-1, CALCA, CTN (CALCITONIN), UCN*, or *UTS2* under the control of the U6 promoter (n = 4). For *VIP*, HEK293T cells overexpressing *VIP* were used, since endogenous *VIP* expression in the MNs was very low.
**e,** Neurite lengths of control MNs transduced with lentiviruses expressing shRNAs targeting *AR*, *CALCA, CGRP-1, CTN, UCN, UTS2*, or *VIP* (n = 3 for shRNAs targeting *AR* and neuropeptides; n = 9 for control shRNA). No significant differences were detected.
**f,** Time-course analysis of the expression of phosphorylated signaling proteins in control MNs treated with each

neuropeptide determined by Western blotting (related to Fig. 6h).
**g,** Quantitative RT–PCR analysis of neuropeptide receptors in control and SBMA MNs after 4 weeks of monolayer culture (n = 4). UTS2R, UTS2 receptor; CLR and RAMP1, CGRP-1 receptors; CRHR1 and CRHR2, UCN receptors; and VIPR1 and VIPR2, VIP receptors.
**h,** Neurite length (left), cell viability (MTS assay, middle), and cytotoxicity (LDH assay, right) of control MNs treated for 1 week with 0.0001% DMSO or signaling inhibitors (100 nM SP600125, a JNK inhibitor, or 100 nM U0126, a MEK inhibitor) after 3–4 weeks of monolayer culture (n = 3). No significant differences were detected.
**i,** Neurite length (left), cell viability (MTS assay, middle), and cytotoxicity (LDH assay, right) of control MNs treated for 1 week with 0.0001% DMSO or UTS2 receptor inhibitors (10 nM SB657510 or 1 nM GSK1562590) after 4 weeks of monolayer culture (n = 3). No significant differences were detected.
Data are presented as the mean ± SEM; *, $p < 0.05$; **, $p < 0.01$; Student's *t* test or one-way ANOVA followed by Dunnett's multiple comparisons test.
