## Supplementary_Figure_Data for "Early pathogenesis of spinal and bulbar muscular atrophy uncovered by human iPSC-derived motor neurons highlights pathogenic neuropeptides as therapeutic targets"

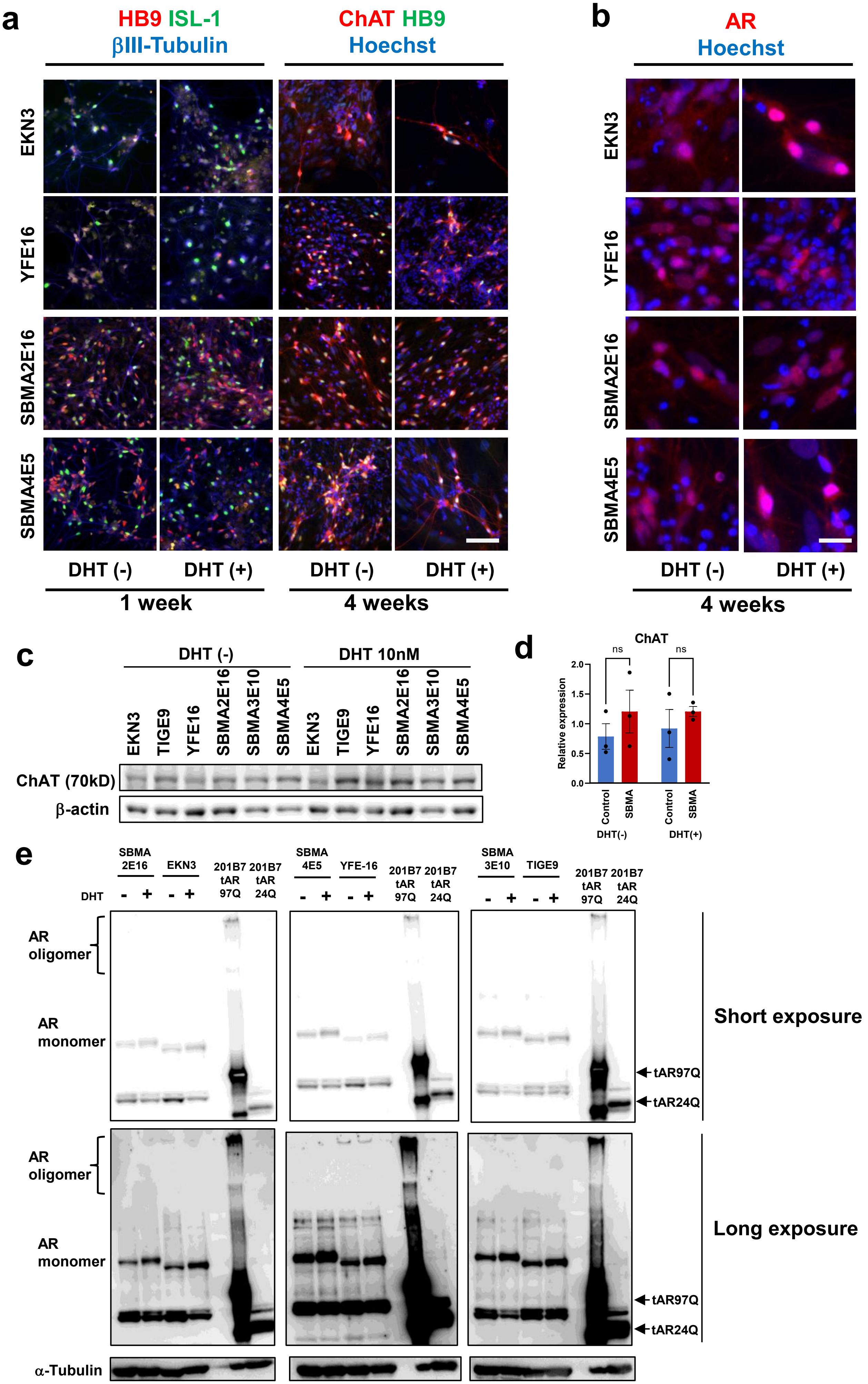


Supplementary Fig. 1 Onodera et al.

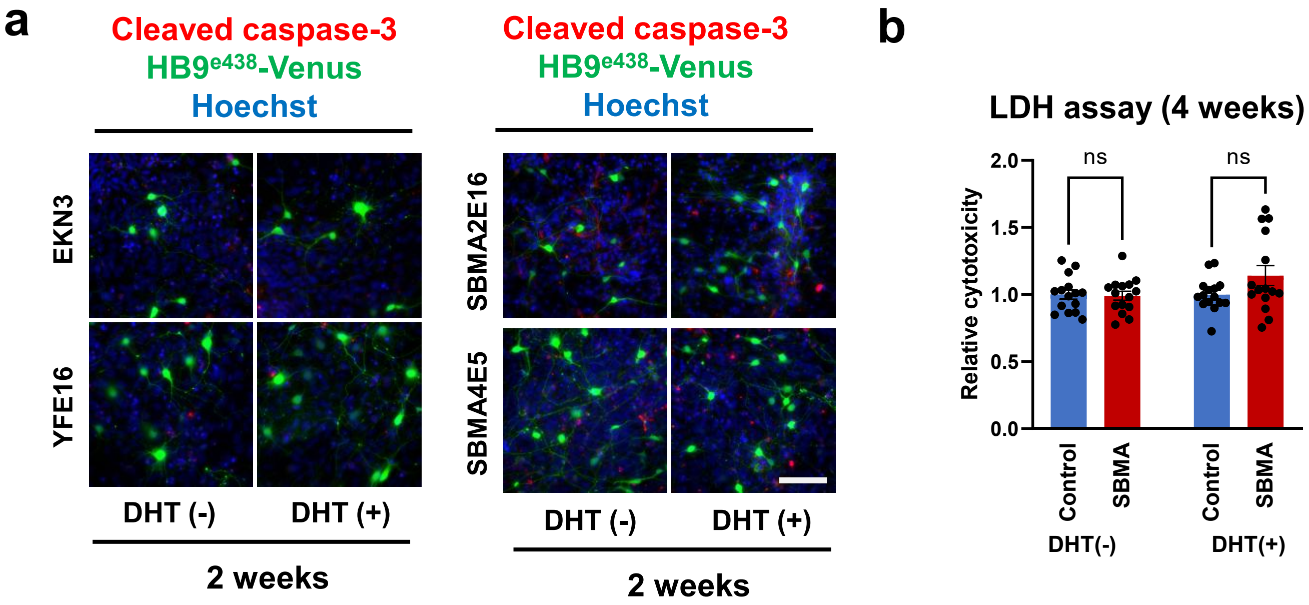


Supplementary Fig. 2 Onodera et al.

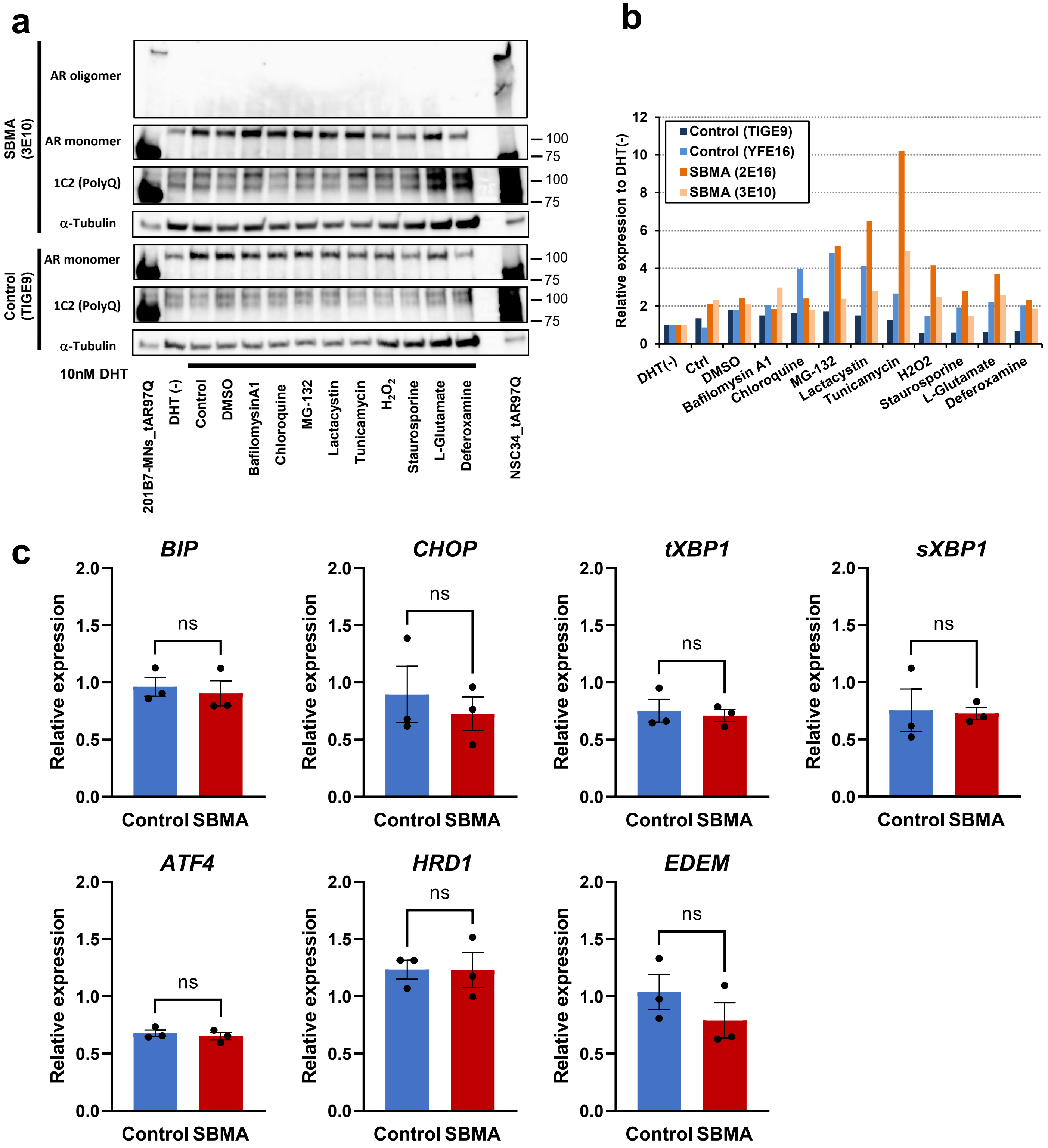


**Supplementary Fig. 3 Onodera et al.**

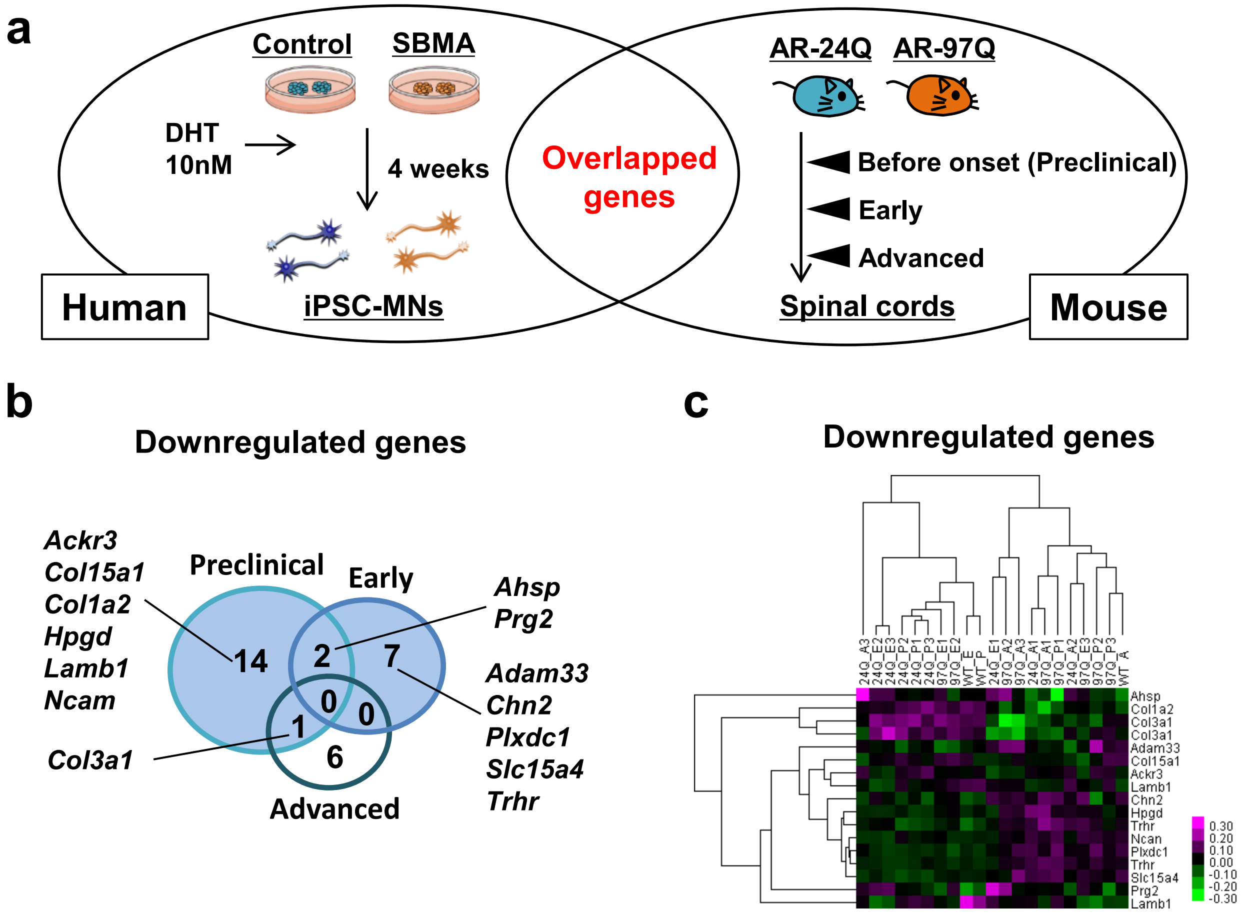


Supplementary Fig. 4 Onodera et al.

a

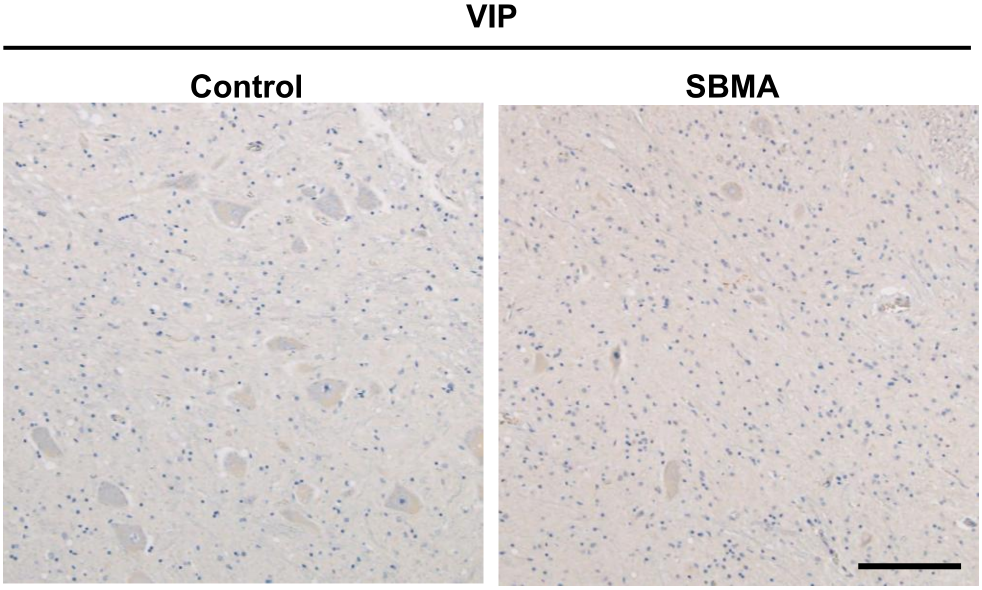


Supplementary Fig. 5 Onodera at al.

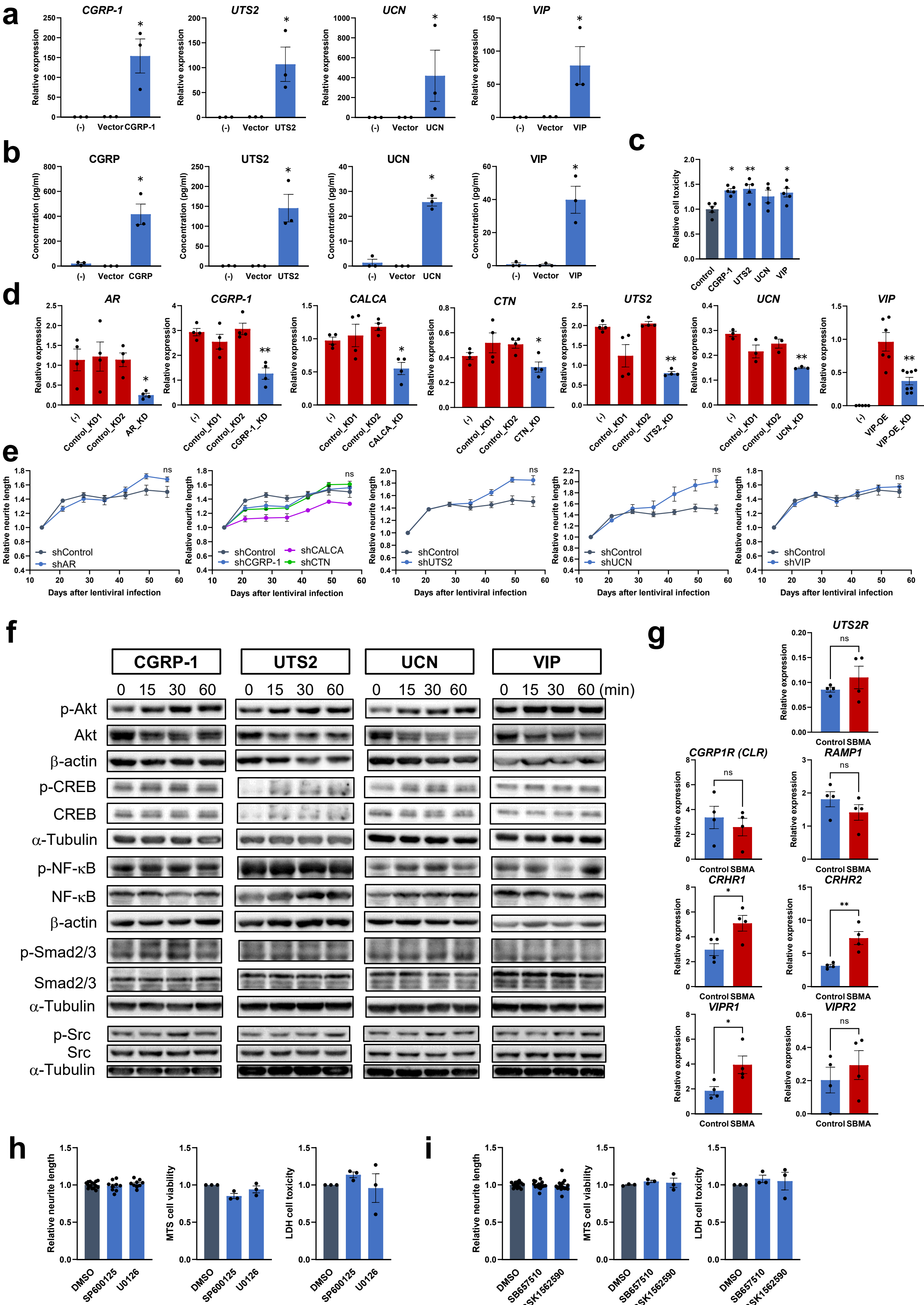


Supplementary Fig. 6 Onodera et al.
